# Unraveling the phylogenetic signal of gene expression from single-cell RNA-seq data

**DOI:** 10.1101/2024.04.17.589871

**Authors:** Joao M Alves, Laura Tomás, David Posada

## Abstract

Single-cell RNA sequencing (scRNA-seq) has transformed our understanding of phenotypic heterogeneity. Although the predominant focus of scRNA-seq analyses has been assessing gene expression changes, several approaches have been proposed in recent years to identify changes at the DNA level from scRNA-seq data. In this study, we evaluated the relative performance of six strategies for calling single-nucleotide variants from scRNA-seq data using 381 single-cell transcriptomes from five cancer patients. Specifically, we focused on the quality of the inferred genotypes and the resulting single-cell phylogenies. We found that scAllele, Monopogen, and Monovar consistently returned phylogenetically informative genotype calls, providing more precise signals of discrimination between tumor and normal cells within heterogeneous samples and among distinct subclonal lineages in longitudinal samples. In addition, we evaluated the evolution of gene expression along the cell phylogenies. While most transcriptomic variation was very plastic and did not correlate with the cell phylogeny, a group of genes associated with cell cycle processes showed a strong phylogenetic signal in one of the patients, underscoring a potential link between gene expression patterns and lineage-specific traits in the context of cancer progression. In summary, our study highlights the potential of scRNA-seq data for inferring cell phylogenies to decipher the evolutionary dynamics of cell populations.

## Introduction

Single-cell RNA sequencing (scRNA-seq) is a powerful technology that allows researchers to explore the gene expression profiles of individual cells, providing insights that are impossible to obtain through traditional RNA sequencing methods (Heumos et al. 2023; Jovic et al. 2022; Lim, Lin, and Navin 2020). While the majority of scRNA-seq analyses have focused on uncovering gene expression differences within complex tissues, recent efforts have emerged to identify DNA changes from scRNA-seq data (F. Liu et al. 2019; Quinones-Valdez et al. 2022; Dou et al. 2023; Muyas et al. 2023; X. Liu et al. 2023). This broadening of scRNA-seq’s applicability, leveraging the wealth of existing datasets, has the potential to reveal the genetic variation that drives cellular heterogeneity and contributes to disease pathogenesis (García-Nieto, Morrison, and Fraser 2019).

Nevertheless, several challenges complicate the detection of genomic variants from scRNA-seq data. Beyond the inherent issues of low coverage and high drop-out rates typical in scRNA-seq (Ziegenhain et al. 2017), factors such as allele-specific expression and alternative splicing may also introduce significant uncertainty in the genotype calls (Kim et al. 2015). Consequently, subsequent inferences based on these calls might be influenced by their quality.

In this study, we evaluate the performance of distinct variant calling strategies for identifying single-nucleotide variants (SNVs) from scRNA-seq data and their impact on downstream evolutionary inferences. The methods assessed include scAllele (Quinones-Valdez et al. 2022), SCOMATIC (Muyas et al. 2023), Monopogen (Dou et al. 2023), and Phylinsic (X. Liu et al. 2023), specifically tailored for scRNA-seq data, as well as HaplotypeCaller (Poplin et al. 2017) and Monovar (Zafar et al. 2016), initially designed for bulk and single-cell DNA sequencing data, respectively.

To showcase a compelling application of scRNA-seq genotypes in the context of cancer evolution, we explored the relationship between genetic changes along cell lineages and gene expression heterogeneity at the single-cell level. Our findings offer a glimpse into the potential of scRNA-seq data for simultaneously unraveling the complex landscape of cancer genomics and transcriptional heterogeneity.

## Material & Methods

### Data retrieval

We retrieved 381 scRNA-seq datasets from the Sequence Read Archive (SRA), corresponding to 22, 85, and 102 cells from three breast cancer (BC) patients (BC01, BC03, BC07) (Chung et al. 2017) and 45 and 127 cells from two multiple myeloma (MM) patients (MM16, MM34) (Fan et al. 2018). All the datasets except for MM34 included healthy and tumor cells. MM34 is a treatment-refractory patient with a bone marrow (BM) sample collected at the time of diagnosis and an extramedullar (EM) sample collected after two months of treatment. Cells in the BM sample are labeled as BM-like and EM-like cells. We also downloaded whole-exome sequencing data from healthy (hbWES) and tumor (tbWES) bulk samples for the BC patients. A list of the individual datasets and corresponding accession codes is available in **Table S1**.

### Read alignment

For the scRNA-seq datasets, we aligned the raw reads to the human reference genome GRCh38 using STAR (Dobin et al. 2013) (v2.7.9a). Following the GATK best practices for RNA-seq data (https://gatk.broadinstitute.org/hc/en-us/articles/360035531192-RNAseq-short-variant-discovery-SNPs-Indels), we marked PCR duplicates using Picard MarkDuplicates (v2.25.5), split the reads into individual exon segments using GATK SplitNCigarReads (v4.2.6.1), and performed base quality score recalibration with GATK BaseRecalibrator and GATK ApplyBQSR.

For the bulk WES datasets, we aligned the raw reads to the human reference genome GRCh38 using BWA-*mem* (H. Li and Durbin 2009). After mapping, we followed the GATK data pre-processing workflow (https://gatk.broadinstitute.org/hc/en-us/articles/360035535912-Data-pre-processing-for-variant-discovery), which involved marking PCR duplicates using Picard MarkDuplicates, and performing base quality score recalibration with GATK BaseRecalibrator and GATK ApplyBQSR.

### Variant calling

We called somatic single-nucleotide variants (SNVs) using six different strategies detailed below. scAllele (Quinones-Valdez et al. 2022), SCOMATIC (Muyas et al. 2023), Monopogen (Dou et al. 2023), and Phylinsic (X. Liu et al. 2023) were specifically developed for scRNA-seq data, while GATK HaplotypeCaller (Poplin et al. 2017) and Monovar (Zafar et al. 2016) were designed for bulk- and single-cell DNA-seq data, respectively. To ensure a fair comparison, we ran all tools under default settings and, when possible, applied the same filtering strategy for the resulting calls.

We only considered biallelic SNVs. We defined 0 as the reference allele and 1 as the alternative allele. Accordingly, the possible genotypes were homozygous for the reference allele (0/0), heterozygous (0/1), and homozygous for the alternative allele (1/1).

### HaplotypeCaller

We ran GATK HaplotypeCaller in multisample mode following the authors’ guidelines. First, we generated GVCF files for each BAM file (one per cell) using the “-ERC GVCF” option, followed by CombineGVCFs and GenotypeGVCFs to merge all variant calls from the same patient into a single VCF. The full command lines were as follows:

> *gatk HaplotypeCaller -R hg38*.*fa -I $*{*sample*}.*bam -O $*{*sample*}.*g*.*vcf*.*gz -ERC GVCF*
>
> *gatk CombineGVCFs -R hg38*.*fa --variant $*{*sample1*}.*g*.*vcf*.*gz --variant $*{*sample2*}.*g*.*vcf*.*gz --variant (*…*) -O $*{*patient*}*-HC*.*g*.*vcf*.*gz*
>
> *gatk GenotypeGVCFs -R hg38*.*fasta -V $*{*patient*}*-HC*.*g*.*vcf*.*gz -O $*{*patient*}*-HC*.*vcf*.*gz*

We only kept biallelic SNVs displaying a “PASS” flag. For the breast cancer patients, we removed germline single-nucleotide polymorphisms (SNPs) identified in the bulk samples (see below). In addition, we set as missing data any genotype supported by less than five reads and 0/1 and 1/1 genotypes with less than two reads for the alternative allele.

### Monovar

We ran Monovar using the authors’ recommended settings (see https://github.com/KChen-lab/MonoVar). We converted all the single-cell BAM files from the same patient into a single mpileup with samtools mpileup and used it as input for Monovar. The command line was as follows:

> *samtools mpileup -BQ0 -d10000 -f hg38*.*fa -q 40 -b $*{*patient_mpileup*}.*Path* | *monovar*.*py -p 0*.*002 -a 0*.*2 -t 0*.*05 -f hg38*.*fa -b $*{*patient_mpileup*}.*Path -o $*{*patient*}*-Monovar*.*vcf*

Afterward, we filtered the resulting calls using the same strategy described above for HaplotypeCaller.

### scAllele

For each patient, we ran scAllele under default settings using the following command line:

> *scAllele -b $*{*list_of_BAM_files*} *-o $*{*patient*}*-scAllele*.*vcf -g hg38*.*fa*

Importantly, scAllele does not output genotype calls, but allele count estimates for each SNV site. We thus followed the author’s suggestions and genotyped all cells, considering the number of reads for the reference (REF) and alternative (ALT) alleles plus their sum (DP). For a given cell and SNV, if DP ≥ 5 and ALT = 0, the genotype was set as 0/0. If DP ≥ 5, REF > 0, and ALT ≥ 2, the genotype was set as 0/1. If DP ≥ 5 and REF = 0, the genotype was 1/1. Otherwise, the genotype was set as missing. As before, we removed non-biallelic SNVs. In addition, we removed the germline variants for the breast cancer patients.

### SCOMATIC

Since SCOMATIC requires barcoded BAM files, we added a “CB” tag (cell-barcode tag) to each single-cell BAM using appendCB (https://github.com/ruqianl/appendCB). Then, we merged the individual BAM files into a single BAM file with reads from healthy and tumor cells for each patient. Following the SCOMATIC guidelines (https://github.com/cortes-ciriano-lab/SComatic), we ran the *BaseCellCounter*.*py* script to generate the read counts. We then used the *MergeBaseCellCounts*.*py* script to merge the read counts for all cells for each patient. Next, we ran the *BaseCellCalling*.*py* script to obtain the variant calls. As recommended by the authors, we applied the *variant intersection* command to remove SNVs located in “low-quality” regions of the human genome (e.g., repeats or regions of low complexity). The command lines used were:

> ^*#*^*base cell counter*
>
> *python3*.*7 BaseCellCounter*.*py --bam $*{*sample*}*-merged*.*bam --ref hg38*.*fa --chrom all --out_folder*
>
> *$*{*path*} *--min_bq 10 --min_dp 2 --tmp_dir $*{*path_temp*} *--min_cc 1*
>
> ^*#*^*merge cell counts*
>
> *python3*.*7 MergeBaseCellCounts*.*py --tsv_folder $*{*path*} *--outfile $*{*patient*}*-Fullset*.*tsv*
>
> ^*#*^*variant calling step 1*
>
> *python3*.*7 BaseCellCalling*.*step1*.*py --infile $*{*patient*}*-Fullset*.*tsv --outfile $*{*patient*}*-Fullset --ref hg38*.*fa --min_cells 2 --min_ac_cells 1 --min_ac_reads 1 --min_cell_types 1*
>
> ^*#*^*variant calling step 2*
>
> *python3*.*7 BaseCellCalling*.*step2*.*py --infile $*{*patient*}*-Fullset*.*calling*.*step1*.*tsv --outfile*
>
> *$*{*patient*}*-Fullset --editing RNAediting/AllEditingSites*.*hg38*.*txt --pon PoNs/PoN*.*scRNAseq*.*hg38*.*tsv*
>
> ^*#*^*remove variable sites in low-quality regions of the human genome:*
>
> *bedtools intersect -header -a $*{*patient*}*-Fullset*.*calling*.*step2*.*tsv -b UCSC*.*k100_umap*.*without*.*repeatmasker*.*bed* | *awk ‘$1 ∽ /∧#/* || *$6 “PASS”‘ >*
>
> *$*{*SAMPLE*}*-Fullset*.*calling*.*step2*.*pass*.*tsv*

Finally, we used the *SingleCellGenotype*.*py* script to extract the read counts for each cell and site. Considering that SCOMATIC only outputs the read counts for either the reference or the alternative allele, we could not apply the same filters described before. In this case, we set the genotypes to missing data if the site had less than two reported reads, otherwise to homozygous reference if reads were for the reference allele, and to heterozygous if reads were for the alternative allele. Note that in this case, we could not call homozygous alternative genotypes.

### Monopogen

We ran Monopogen following the recommended workflow (https://github.com/KChen-lab/Monopogen). Using the “CB-tagged” BAM files generated for SCOMATIC for each patient (which contains the aligned reads from all healthy and tumor cells), we ran the *preProcess* script to remove reads with more than three alignment mismatches. Afterward, we called *germline* SNPs for each chromosome with the corresponding 1000 Genomes imputation panel (https://ftp.1000genomes.ebi.ac.uk/vol1/ftp/data_collections/1000G_2504_high_coverage/working/20201028_3202_phased/). Finally, we ran a three-step procedure to identify *somatic* SNVs and to obtain the set of genotypes across cells. The full workflow was set as follows:

> ^*#*^*pre-process step*
>
> *Monopogen*.*py preProcess -b $*{*patient_bam*}.*lst -o $*{*patient*} *-a $*{*monopogen_apps*}
>
> ^*#*^*SNP calling*
>
> *Monopogen*.*py germline -s all -o $*{*patient*} *-g hg38*.*fa -a $*{*monopogen_apps*} *-p*
>
> *$*{*phase_panel_chr*} *-r $*{*chr*}.*lst*
>
> ^*#*^*SNV calling*
>
> *Monopogen*.*py somatic -a $*{*monopogen_apps*} *-i $*{*patient*} *-r $*{*chr*}.*lst -l $*{*patient_samples*}.*csv -s featureInfo -g hg38*.*fa*
>
> *Monopogen*.*py somatic -a $*{*monopogen_apps*} *-i $*{*patient*} *-r $*{*chr*}.*lst -l $*{*patient_samples*}.*csv -s cellScan -g hg38*.*fa*
>
> *Monopogen*.*py somatic -a $*{*monopogen_apps*} *-r $*{*chr*}.*lst -i $*{*patient*} *-l $*{*patient_samples*}.*csv -s LDrefinement -g hg38*.*fa*

For downstream analysis, we kept only SNVs showing an “LDrefine_merged_score > 0.25” (a metric used to distinguish somatic from germline variants) and an alternative allele frequency above 0.3. Since Monopogen already removes germline variants and performs genotype imputation for all cells and variable sites, we did not apply additional filters.

### Phylinsic

Although Phylinsic was primarily designed to work with 10X Genomics scRNA-seq data, we followed the author’s suggestions to adjust the snakefile to Smart-seq2 data (https://github.com/U54Bioinformatics/PhylinSic_Project/issues/1). Cell categories were set as “healthy” or “tumor” –except for patient MM34, where we followed the labels described in the original publication. To execute the full Phylinsic pipeline, we used the following command line:

> *snakemake --snakefile updated_snakefile*.*txt*

Like Monopogen, Phylinsic already filters out germline variants and performs genotype imputation; hence, we did not apply additional filters.

### Bulk WES variant calling

We also identified germline and somatic mutations in the bulk WES samples. For the germline mutations, we ran GATK HaplotypeCaller on the healthy bulk samples. Afterward, we ran the Variant Quality Score Recalibration (*VQSR*) pipeline to remove low-quality variants. We used the paired healthy/tumor variant calling approach implemented in the MuTect2 software to identify somatic mutations. The output somatic callsets were then filtered using the *FilterMutectCalls*.

### Evaluation of the scRNA-seq variant calls

In order to evaluate the relative reliability of the scRNA-seq genotypes, we computed four different measures for each scRNA-seq variant calling strategy, two at the population level and two at the single-cell level:

*SNV patient recall* is the proportion of SNVs inferred in the tbWES datasets also identified in the scRNA-seq data from the same patient.

*SNP patient recall* is the proportion of SNPs inferred in the hbWES datasets that were also identified in the scRNA-seq data from the same patient. This measure was only calculated for the three methods that do not directly filter out germline variants - HaplotypeCaller, Monovar, and scAllele.

*SNP cell recall* is, for a given cell, the proportion of SNPs in the hbWES datasets also identified in the scRNA-seq data.

*Allelic dropout rate* is, for a given cell, the proportion of scRNA-seq genotypes at hetSNP sites in the hbWES dataset that were identified as homozygous (0/0 or 1/1).

### Cell phylogeny estimation

It is relatively well-established that missing data can compromise phylogenetic accuracy (Smith et al. 2020). As such, prior to the phylogenetic reconstruction, we removed sites displaying more than 75% missing genotypes for either healthy or tumor cells and cells with more than 75% missing genotypes for any of the scRNA-seq variant calling strategies. As a consequence, some datasets experienced a reduction. The final number of cells was 22, 59, 88, 39, and 113 for BC01, BC03, BC07, MM16, and MM34, respectively.

We used CellPhy (Kozlov et al. 2022) to reconstruct the cell phylogenies using maximum likelihood under the GT16 model with 100 bootstrap replicates of the genotype matrix. Branch support estimates were obtained using the transfer bootstrap expectation metric (Lemoine et al. 2018). The command line for all runs was:

> *cellphy*.*sh RAXML --all --msa $*{*set*}.*vcf --model GT16+FO+E --msa-format auto --prefix $*{*set*} *--bs-tree autoMRE*{*100*} *--bs-metric tbe --prob-msa off*.

### Reliability of the cell phylogeny

The actual cell phylogeny is unknown, but we would like to have some measure of the reliability of the inferred cell trees. For that purpose, we employed two different strategies that focus on how well the trees differentiate among healthy and tumor cells, as we expect tumor cells to form an exclusive cluster with a single origin (in phylogenetic terms, a monophyletic group, or clade). First, we measured the mean phylogenetic distance between cells belonging to the same class (i.e., healthy vs. tumor) using the “distTips” function from the adephylo R package (Jombart, Balloux, and Dray 2010). In parallel, we computed Moran’s I index (Moran 1950) with the R package “PATH” (Schiffman et al. 2022) to assess the phylogenetic autocorrelation within healthy and tumor cells. We computed both measures based on the 100 bootstrap trees previously generated with CellPhy to account for the uncertainty in the cell trees.

### Phylogenetic signal of gene expression

Cell phylogenies derived from scRNA-seq data enable the assessment of how a cell’s gene expression levels are statistically dependent on its evolutionary history, providing insights into the “phylogenetic signal” embedded within gene expression patterns. Here, we measured the phylogenetic signal of gene expression in the tumor cells with Pagel’s lambda (Pagel 1999), using the function “tree_physig” from the *sensiPhy* R package (Paterno, Penone, and Werner 2018) and the cell phylogenies obtained with the scAllele genotypes. Again, to incorporate the phylogenetic uncertainty of the cell tree estimates in the analysis, we computed *lambda* across the 100 bootstrap trees previously generated with CellPhy. To measure the gene expression levels, we obtained read counts for all cells from the NCBI Gene Expression Omnibus database (accession codes available in **Table S1**) and normalized them using the *DESeq2* R package (Love, Huber, and Anders 2014). For each patient, a principal component analysis (PCA) was used to visualize the patterns of similarity in gene expression between cells (**fig. S1**). Finally, we performed an enrichment analysis for those genes whose expression showed a significant phylogenetic signal (mean lambda > 0.5 and upper-CI p-value < 0.05) using the “gseGO” function of the *clusterProfiler* R package (Wu et al. 2021). The parent terms for each GO were obtained using the “getParents” function in the *GOSim* R package (Fröhlich et al. 2007).

### Variant annotation and mapping

We annotated the scAllele SNV calls from the MM34 patient using Annovar (Wang, Li, and Hakonarson 2010) and mapped all non-synonymous SNVs to the corresponding cell tree using the *mutmap* function in CellPhy (Kozlov et al. 2022).

## Results

### Different strategies result in distinct scRNA-seq genotype calls

The number of somatic SNVs identified varied considerably depending on the strategy used to call them. Across patients, Monovar and HaplotypeCaller consistently identified the largest amount, with counts being, in most cases, one order of magnitude higher than for the other strategies specifically designed for scRNA-seq data (**fig. 1A**). In contrast, SCOMATIC returned the lowest number of SNVs across all sets evaluated; for the multiple myeloma patients, it only identified four SNVs when other methods identified hundreds to thousands of SNVs.

**Figure 1.**
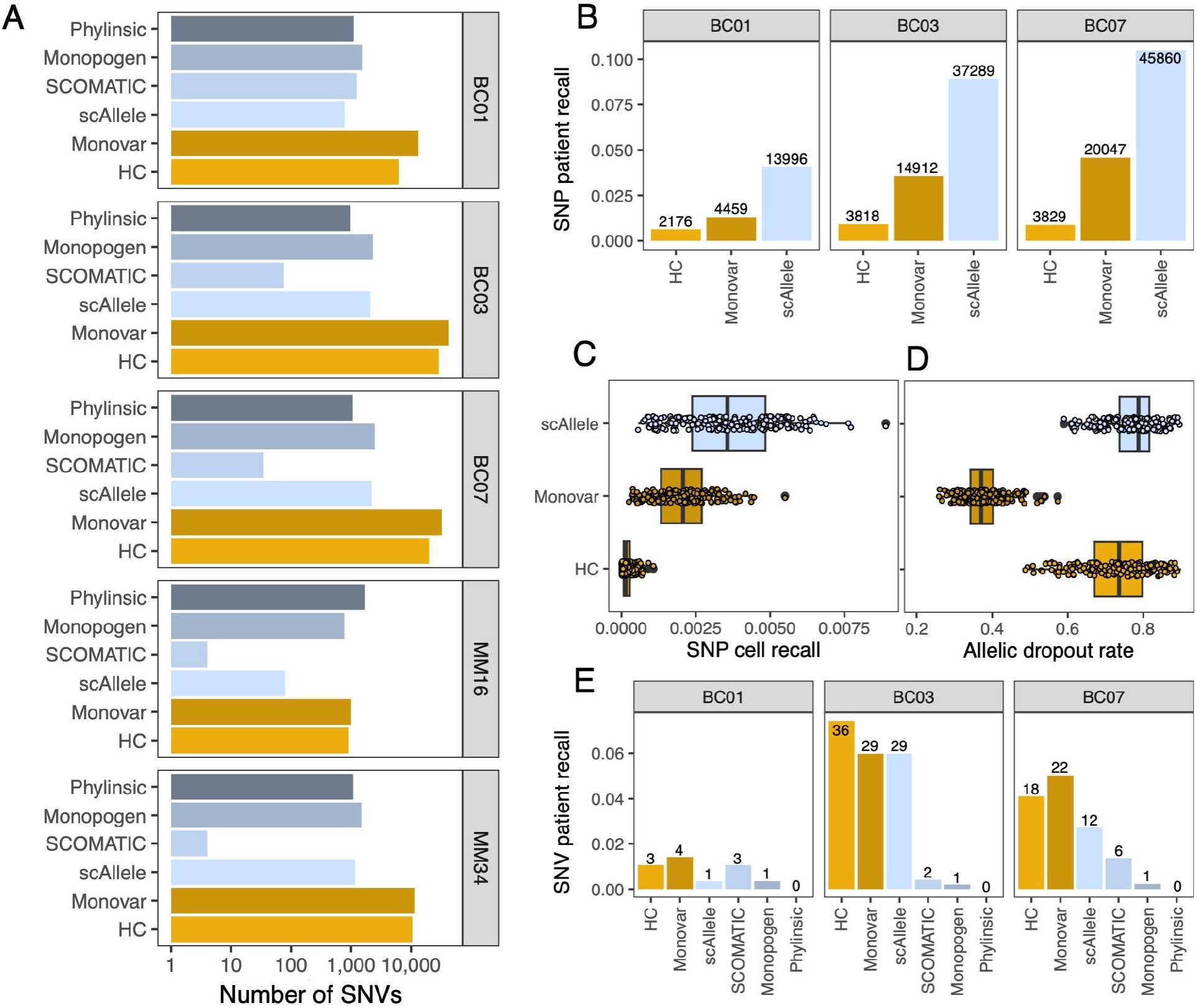
Variant calling from scRNA-seq data. **A**. Barplots depicting the number of SNVs called by the different strategies across breast cancer (BC) and multiple myeloma (MM) patients. Note the log scale in the x-axis. **B**. SNP patient recall for the BC patients. Only HaplotypeCaller (HC), Monovar, and scAllele are included here, as all other methods do not call germline variants. The number of SNPs is shown above each bar **C**. SNP cell recall for the BC patients combined. **D**. Allelic dropout rate for the BC patients combined. **E**. SNV patient recall for BC patients. The number of SNVs is shown above each bar.

### SNP and SNV recall rates are low; allelic dropout is rampant

Germline SNPs are expected to be present in all cells from a given patient, making them particularly valuable for evaluating scRNA-seq variant calling strategies. We computed SNP recall by comparing the scRNA-seq calls - obtained with the three variant calling methods that do not distinguish germline from somatic variants (HaplotypeCaller, Monovar, and scAllele) - with the SNP calls from the bulk healthy WES (bhWES) data. When considering the calls made for a given patient, all three methods showed low SNP recall rates, always below 11%. scAllele consistently identified the highest proportion of SNPs, corresponding to ∽14,000 - 46,000 SNPs (**fig. 1B**). At the individual cell level, we observed minimal SNP recall rates, that varied notably from cell to cell, with scAllele still showing the largest values (**fig. 1C**). Interestingly, the variant allele frequency distributions obtained with scAllele were quite similar to those derived from the bhWES data, while this was not the case for Monovar and, mainly, HaplotypeCaller (**fig. S2**). The observed allelic dropout (ADO) rates were particularly high for scAllele (ADO ∽ 0.8) and HaplotypeCaller (ADO ∽ 0.75), but half of that for Monovar (ADO ∽ 0.37) (**fig. 1D**). However, it is worth noting that Monovar tends to genotype variable sites as heterozygous (i.e., 0/1) even when reads are only observed for the alternative allele, which could partly explain this result. Finally, only a very small fraction (0-8%) of the SNVs identified in the bulk tumor WES (btWES) data were also detected in the corresponding scRNA-seq datasets, with HaplotypeCaller, Monovar, and scAllele returning the largest numbers, in the scale of dozens (**fig. 1E**).

### The inferred cell phylogeny depends on the scRNA-seq variant calling approach

Different strategies for obtaining SNV genotypes from scRNA-seq data result in distinct cell phylogenies (**fig. 2**). In the case of HaplotypeCaller, for instance, the inferred trees exhibit long terminal branches and relatively short internal ones. While this pattern is expected in the phylogenies of cells sampled from growing populations, most HaplotypeCaller trees display healthy cells intermixed with tumor cells, a result that seems unrealistic. Similar patterns were observed for SCOMATIC where, despite a strong bootstrap support for many nodes, the trees contain many polytomies (i.e., unsolved relationships), and tumor cells often fail to cluster together. In contrast, the trees derived from the scAllele, Monovar, and Monopogen genotypes showed tumor cells predominantly grouping with high support. Finally, Phylinsic calls resulted in trees with incomplete separation between healthy and tumor cells.

**Figure 2.**
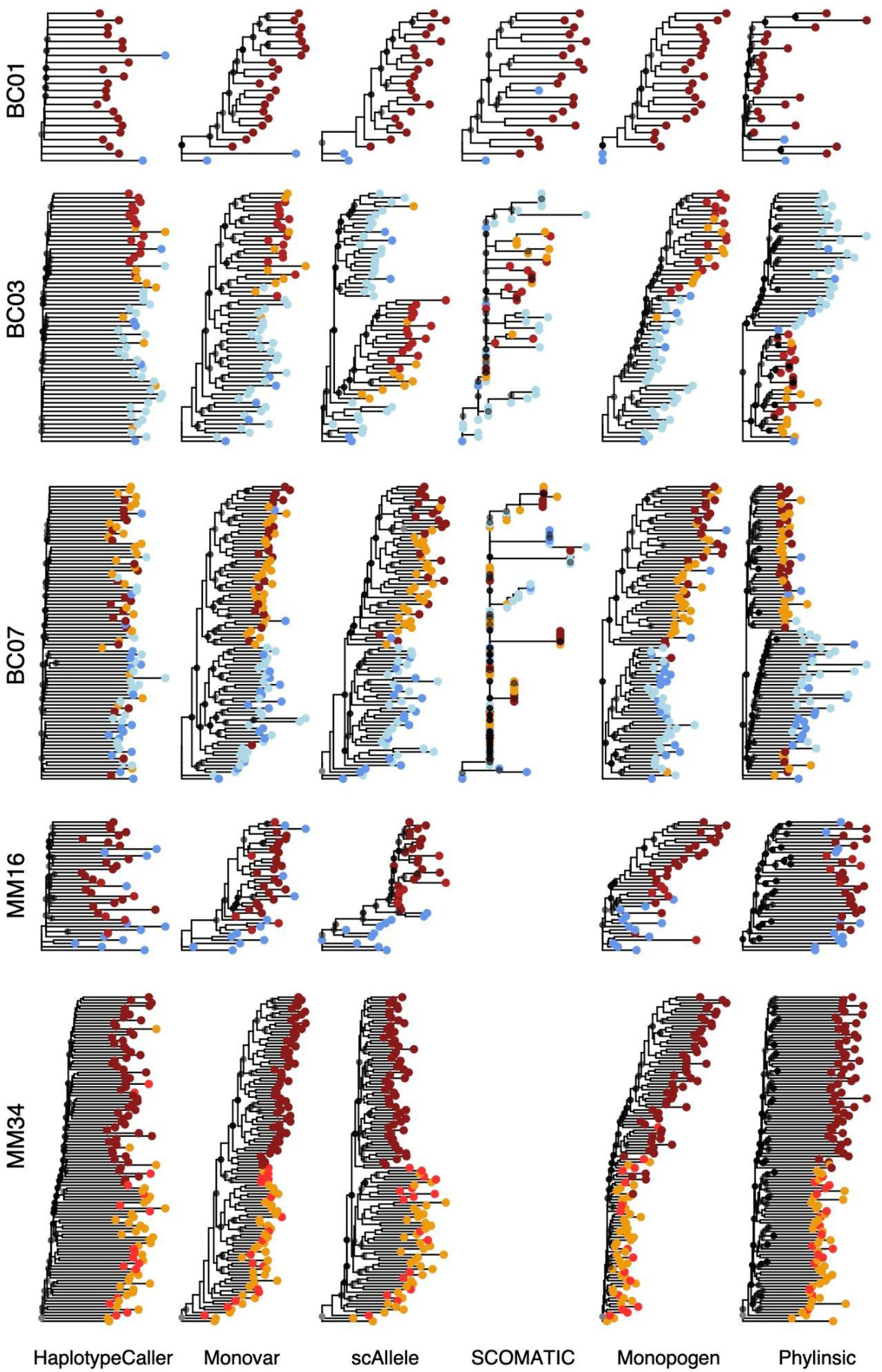
Cell phylogenies inferred from scRNA-seq data. CellPhy trees for all patients according to different scRNA-seq variant calling approaches. For display purposes, all trees were rooted within each patient using the same arbitrary cell as outgroup. For BC01, BC03, BC07, and MM16, the colored tips represent different cell categories or sampling locations: healthy cells - blue; primary tumor cells - dark red; lymph-node tumor cells - dark orange. For MM34, tip colors represent different cell categories: EM: dark red; EM-like: light red; BM-like: dark orange. Only bootstrap values above 50% are depicted, using a continuous transparency scale where solid black circles represent a 100% value. Because SCOMATIC only detected four SNVs for the MM patients, the cell phylogenies were not estimated in this case.

In the case of patient MM34, most cell phylogenies provide a fairly clear separation between cells sampled at different time points (i.e., EM *vs*. BM-like and EM-like cells), strongly supporting a monoclonal extramedullary seeding event. Nevertheless, none of the methods supported two distinct genetic lineages within the bone marrow corresponding to BM-like and EM-like cells.

To better estimate the reliability of these cell trees beyond a visual inspection, we computed the pairwise phylogenetic distance between cells of the same class, i.e., healthy vs. tumor (**fig. 3**). Remarkably, scAllele, Monopogen, and Monovar repeatedly returned the lowest phylogenetic distance among cells of the same type. Phylinsic, on the other hand, resulted in relatively large phylogenetic distances in BC01 and MM16 but smaller ones in BC03 and BC07. Finally, both HaplotypeCaller and SCOMATIC often yielded the highest phylogenetic distances between cells of the same type. We obtained similar results using Moran’s autocorrelation index (**fig. S3**). Across all datasets, the highest autocorrelation scores corresponded again to scAllele, Monopogen, and Monovar.

**Figure 3.**
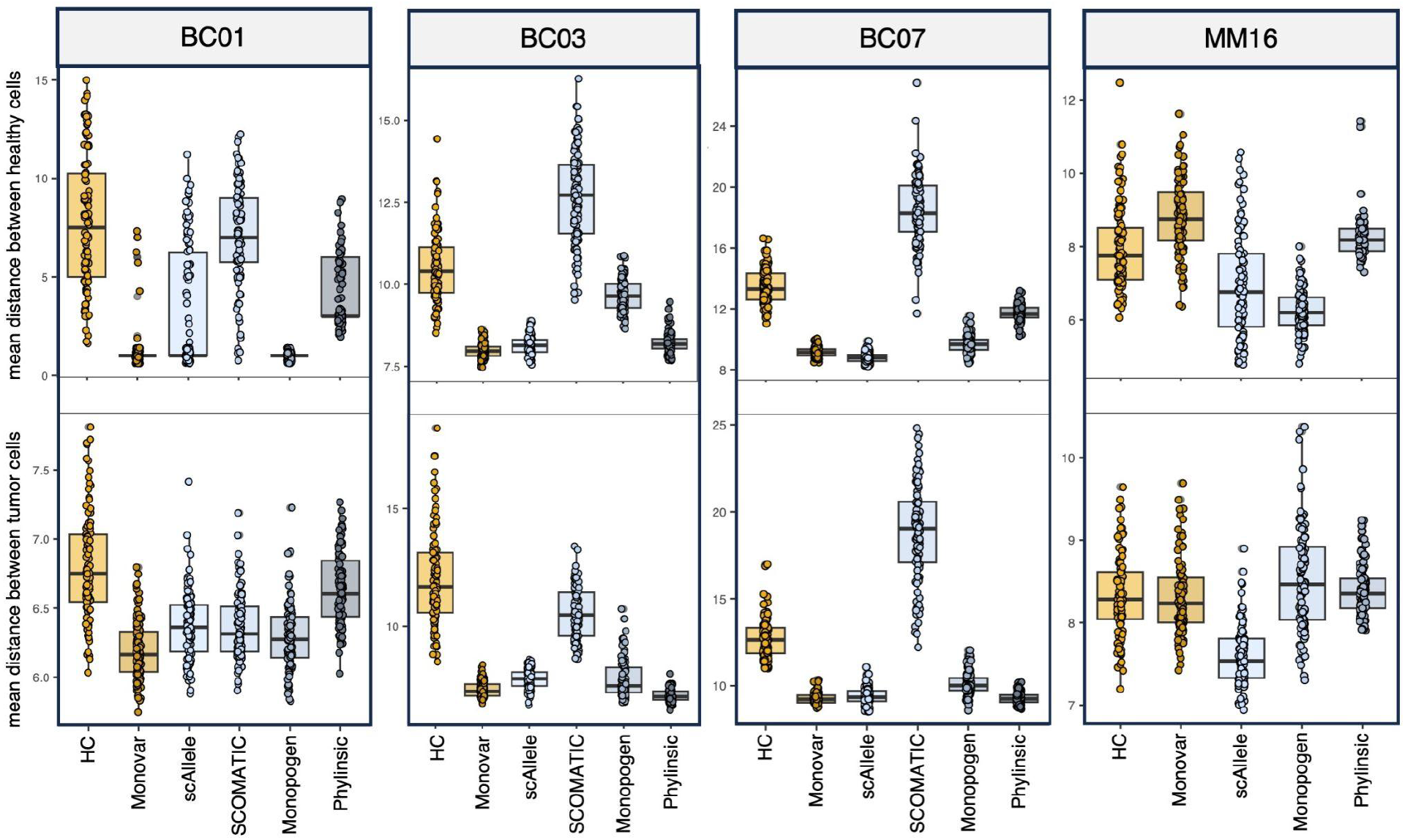
Intra-class phylogenetic distances. Boxplots depicting the phylogenetic distance between cells in the same class (Top=healthy; Bottom=Tumor). Each data point represents the average cell-to-cell phylogenetic distance across 100 bootstrap trees. HC = HaplotypeCaller.

### Strong phylogenetic signal in MM34

Considering its better performance for variant calling and phylogenetic estimation, we estimated the phylogenetic signal for gene expression using only the cell phylogenies derived from the scAllele genotypes. Nevertheless, we obtained similar results when using the genotypes inferred by Monovar (**fig. S4**).

We found a statistically significant phylogenetic signal – measured with the lambda statistic (see Methods) – only in patients BC07 and MM34. In BC07, three genes, *PTPRC, CD69*, and *CYTIP*, displayed significant (p-value < 0.05) mean lambda scores ≥ 0.5, suggesting moderate to high phylogenetic signal (**fig. S5**). Notably, in MM34, 1,877 genes showed a significant mean lambda score ≥ 0.5 (**fig. 4A**). A gene set enrichment analysis of these genes uncovered a significant overrepresentation in cell cycle and chromosome organization processes (**fig. 4B**), suggesting potential alterations in cell division rates between bone marrow (BM-like and EM-like) and extramedullary disseminated (EM) cells. The same genes were mostly down-regulated in the bone marrow without apparent differences in expression levels between the BM-like and the EM-like subgroups (**fig. 4C**). In addition, we identified a non-synonymous mutation in the *NDUFB9* gene along the single branch leading to the EM cell population (**fig. 4C**). Interestingly, although the down-regulation of *NDUFB9* has been linked to increased proliferation and metastatic dissemination in breast cancer (Li et al. 2015), we did not find statistically significant differences in expression levels between the EM and the BM cells (**fig. 4D**).

**Figure 4.**
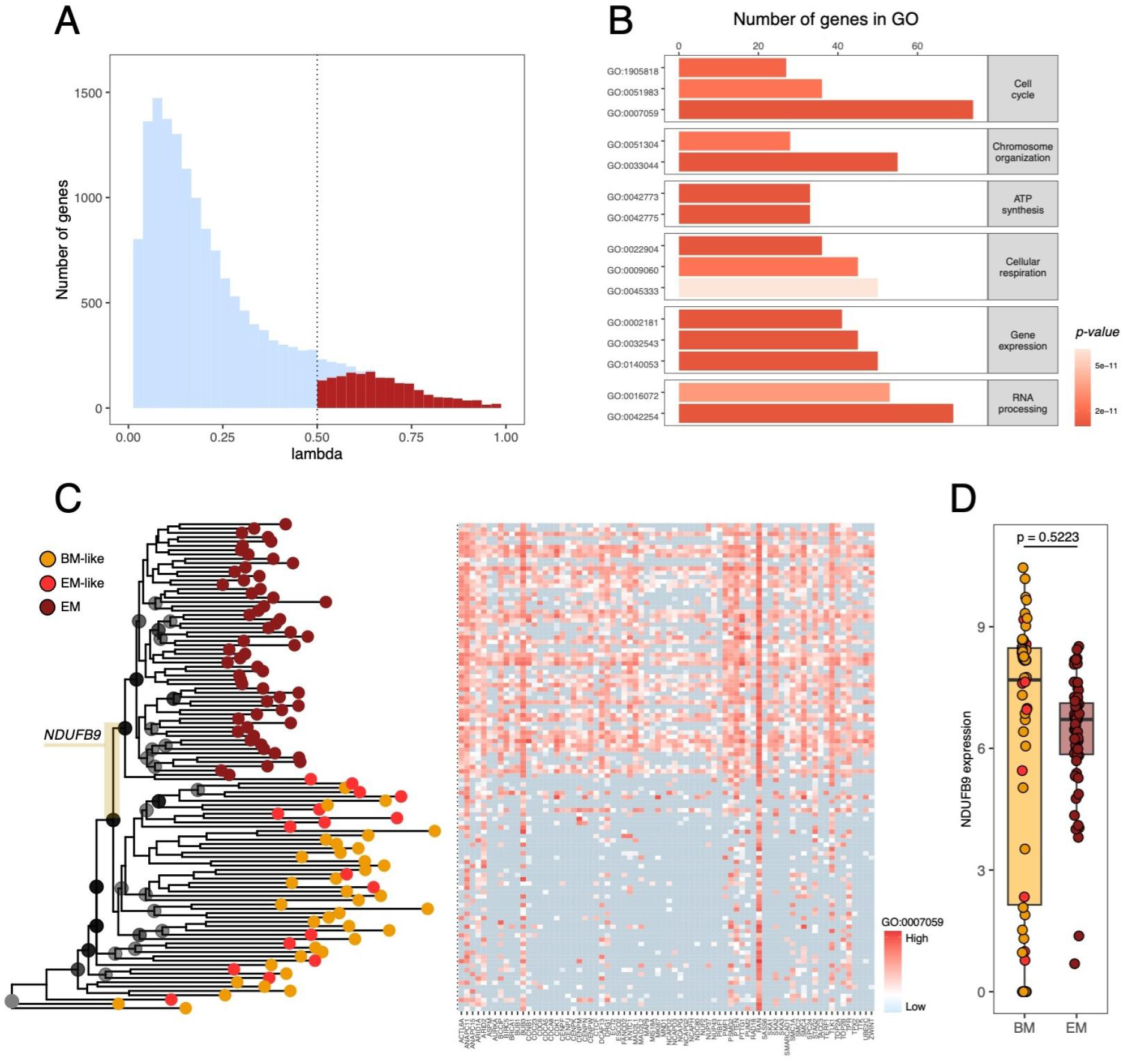
Phylogenetic signal of gene expression in patient MM34. **A**. Mean lambda score for each gene. Genes showing moderate to high phylogenetic signal (i.e., mean lambda score ≥ 0.5 and p-value < 0.05) are colored in red. **B**. Top 15 GO terms in GSEA for genes showing a significant lambda score. **C**. MM34 phylogenetic tree using the scAllele genotypes along with cell-level gene expression profiles for genes comprising GO:0007059. Tip colors represent different cell categories: EM: dark red; EM-like: light red; BM-like: dark orange. Only bootstrap support values equal to or above 50 are indicated, using a continuous transparency scale (with solid black circles representing a branch support of 100%). A non-synonymous mutation in *NDUFB9* is highlighted on the branch leading to the most recent common ancestor of the EM cells. Cells in the tile plot are ordered according to the phylogenetic tree. **D**. Boxplots depicting the expression levels of *NDUFB9* for BM and EM samples. Each data point represents an individual cell, color-coded according to the scheme used in the phylogenetic tree. The t-test *p-value* is shown on top.

## Discussion

scRNA-seq has emerged as a powerful tool in unraveling the intricate phenotypic heterogeneity present within complex tissues. Nonetheless, the value of scRNA-seq data extends beyond understanding cellular diversity. It provides a unique opportunity to identify genomic changes and leverage this information to reconstruct cell phylogenies, ultimately enabling us to trace the evolutionary patterns of gene expression. In this study, we evaluated the potential of six SNV calling strategies to explore the evolutionary history of tumors using scRNA-seq data.

Our findings suggest that some methods result in more reliable evolutionary relationships among cell lineages. Notably, scAllele, Monopogen, and Monovar consistently demonstrated the ability to yield phylogenetically informative genotypes. These methods facilitated the discrimination between tumor and normal cells within heterogeneous samples and enabled the identification of distinct subclonal lineages. For example, the majority of the MM34 scRNA-seq phylogenies suggest a monoclonal extramedullary seeding event, aligning with previous observations derived from copy number profiles (Fan et al. 2018). However, in contrast to the original study, none of the cell phylogenies clustered the bone marrow sampled cells into BM-like and EM-like subgroups. At the same time, we observed that the statistical support for branches within cell categories was relatively low. This limitation in resolving fine-scale evolutionary relationships within subpopulations may stem from the inherent complexities involved in detecting subtle genetic differences at the single-cell level using scRNA-seq data, emphasizing the need for a more cautious interpretation.

A significant benefit of using scRNA-seq data to build cell phylogenies is that it allows us to study the evolution of gene expression along cell lineages. Shared ancestry can lead to phenotypic similarity between related cell lineages, a phenomenon known in evolutionary biology as “phylogenetic signal” (Revell, Harmon, and Collar 2008). Consequently, assessing the phylogenetic signal of gene expression holds considerable potential in identifying mutations that drive phenotypic diversity and uncovering signatures of adaptive evolution. Within patient MM34, a notable proportion of genes associated with cell cycle processes exhibited altered expression levels along the cell phylogeny. Remarkably, these genes were particularly active within a specific lineage, suggesting potential differences in proliferation rate, which may have contributed to the rapid extramedullary dissemination of the disease.

We found a notable discrepancy in the phylogenetic signal of gene expression across datasets, with only MM34 exhibiting a strong phylogenetic signal for many genes. We hypothesize that the well-structured phylogeny within MM34, where cells from different spatial locations and distinct microenvironments form established populations with high phylogenetic support, contributes significantly to this observation. In contrast, cell trees for the remaining patients exhibited intermixed cell populations across sampling locations, potentially limiting the detection of phylogenetic signal. It is important to note that the majority of genes in all datasets did not display a significant phylogenetic signal for expression. This outcome is consistent with previous results and underscores the prevalence of transcriptional plasticity (Househam et al. 2022).

Furthermore, our findings emphasize the need to consider phylogenetic uncertainty when estimating the phylogenetic signal for cellular traits. The limited bootstrap support observed in the scRNA-seq phylogenies can result in considerably unstable lambda scores (**fig. S6**). Therefore, *ad hoc* approaches like the one proposed here should be considered when interpreting these trees. Alternatively, Bayesian approaches offer a natural framework for addressing the challenges posed by phylogenetic uncertainty. While acknowledging the limitation of our analysis, centered on a restricted number of datasets with varying cell counts, it is noteworthy that the overall patterns remained remarkably consistent across these datasets. However, these results may only be partially generalizable, particularly to datasets generated using different technologies, such as 10X genomics, where read coverage is often much lower.

In conclusion, our study highlights the valuable insights that can be gained from scRNA-seq data, shedding light on the genomic diversity within cell populations and facilitating a deeper understanding of the evolutionary changes that occur in gene expression along specific cell lineages. Our results offer a comprehensive benchmark for variant detection and phylogenetic reconstruction using scRNA-seq data, providing a valuable guideline for exploring the evolutionary dynamics of cell populations through the analysis of single-cell transcriptomes.

## Ethics approval and consent to participate

Not applicable.

## Consent for publication

Not applicable.

## Availability of data and materials

A list of the public datasets analyzed is in **Table S1**.

## Authors’ contributions

DP conceived the study, and JMA designed the analyses. JMA retrieved and processed all samples. JMA and LT performed the analyses. All authors read and approved the final manuscript.

## Acknowledgments

We thank the Supercomputation Center of Galicia (CESGA; https://www.cesga.es) for providing all the computational resources needed to carry out this study.

## References

Chung, Woosung, Hye Hyeon Eum, Hae-Ock Lee, Kyung-Min Lee, Han-Byoel Lee, Kyu-Tae Kim, Han Suk Ryu, et al. 2017. “Single-Cell RNA-Seq Enables Comprehensive Tumour and Immune Cell Profiling in Primary Breast Cancer.” Nature Communications 8 (May): 15081.

Dobin, Alexander, Carrie A. Davis, Felix Schlesinger, Jorg Drenkow, Chris Zaleski, Sonali Jha, Philippe Batut, Mark Chaisson, and Thomas R. Gingeras. 2013. “STAR: Ultrafast Universal RNA-Seq Aligner.” Bioinformatics 29 (1): 15–21.

Dou, Jinzhuang, Yukun Tan, Kian Hong Kock, Jun Wang, Xuesen Cheng, Le Min Tan, Kyung Yeon Han, et al. 2023. “Single-Nucleotide Variant Calling in Single-Cell Sequencing Data with Monopogen.” Nature Biotechnology, August. 10.1038/s41587-023-01873-x.

Fan, Jean, Hae-Ock Lee, Soohyun Lee, Da-Eun Ryu, Semin Lee, Catherine Xue, Seok Jin Kim, et al. 2018. “Linking Transcriptional and Genetic Tumor Heterogeneity through Allele Analysis of Single-Cell RNA-Seq Data.” Genome Research 28 (8): 1217–27.

Fröhlich, Holger, Nora Speer, Annemarie Poustka, and Tim Beissbarth. 2007. “GOSim--an R-Package for Computation of Information Theoretic GO Similarities between Terms and Gene Products.” BMC Bioinformatics 8 (May): 166.

García-Nieto Pablo E., Ashby J. Morrison, and Hunter B. Fraser. 2019. “The Somatic Mutation Landscape of the Human Body.” Genome Biology 20 (1): 298.

Heumos, Lukas, Anna C. Schaar, Christopher Lance, Anastasia Litinetskaya, Felix Drost, Luke Zappia, Malte D. Lücken, et al. 2023. “Best Practices for Single-Cell Analysis across Modalities.” Nature Reviews. Genetics 24 (8): 550–72.

Househam, Jacob, Timon Heide, George D. Cresswell, Inmaculada Spiteri, Chris Kimberley, Luis Zapata, Claire Lynn, et al. 2022. “Phenotypic Plasticity and Genetic Control in Colorectal Cancer Evolution.” Nature 611 (7937): 744–53.

Jombart, Thibaut, François Balloux, and Stéphane Dray. 2010. “Adephylo: New Tools for Investigating the Phylogenetic Signal in Biological Traits.” Bioinformatics 26 (15): 1907–9.

Jovic, Dragomirka, Xue Liang, Hua Zeng, Lin Lin, Fengping Xu, and Yonglun Luo. 2022. “Single-Cell RNA Sequencing Technologies and Applications: A Brief Overview.” Clinical and Translational Medicine 12 (3): e694.

Kim, Jong Kyoung, Aleksandra A. Kolodziejczyk, Tomislav Ilicic, Sarah A. Teichmann, and John C. Marioni. 2015. “Characterizing Noise Structure in Single-Cell RNA-Seq Distinguishes Genuine from Technical Stochastic Allelic Expression.” Nature Communications 6 (October): 8687.

Kozlov, Alexey, Joao M. Alves, Alexandros Stamatakis, and David Posada. 2022. “CellPhy: Accurate and Fast Probabilistic Inference of Single-Cell Phylogenies from scDNA-Seq Data.” Genome Biology 23 (1): 37.

Lemoine, F., J-B Domelevo Entfellner, E. Wilkinson, D. Correia, M. Dávila Felipe, T. De Oliveira, and O. Gascuel. 2018. “Renewing Felsenstein’s Phylogenetic Bootstrap in the Era of Big Data.” Nature 556 (7702): 452–56.

Li, Heng, and Richard Durbin. 2009. “Fast and Accurate Short Read Alignment with Burrows-Wheeler Transform.” Bioinformatics 25 (14): 1754–60.

Li, Liang-Dong, He-Fen Sun, Xue-Xiao Liu, Shui-Ping Gao, Hong-Lin Jiang, Xin Hu, and Wei Jin. 2015. “Down-Regulation of NDUFB9 Promotes Breast Cancer Cell Proliferation, Metastasis by Mediating Mitochondrial Metabolism.” PloS One 10 (12): e0144441.

Lim, Bora, Yiyun Lin, and Nicholas Navin. 2020. “Advancing Cancer Research and Medicine with Single-Cell Genomics.” Cancer Cell 37 (4): 456–70.

Liu, Fenglin, Yuanyuan Zhang, Lei Zhang, Ziyi Li, Qiao Fang, Ranran Gao, and Zemin Zhang. 2019. “Systematic Comparative Analysis of Single-Nucleotide Variant Detection Methods from Single-Cell RNA Sequencing Data.” Genome Biology 20 (1): 1–15.

Liu, Xuan, Jason I. Griffiths, Isaac Bishara, Jiayi Liu, Andrea H. Bild, and Jeffrey T. Chang. 2023. “Phylogenetic Inference from Single-Cell RNA-Seq Data.” Scientific Reports 13 (1): 12854.

Love, Michael I., Wolfgang Huber, and Simon Anders. 2014. “Moderated Estimation of Fold Change and Dispersion for RNA-Seq Data with DESeq2.” Genome Biology 15 (12): 550.

Moran, P. A. P. 1950. “Notes on Continuous Stochastic Phenomena.” Biometrika 37 (1-2): 17–23.

Muyas, Francesc, Carolin M. Sauer, Jose Espejo Valle-Inclán, Ruoyan Li, Raheleh Rahbari, Thomas J. Mitchell, Sahand Hormoz, and Isidro Cortés-Ciriano. 2023. “De Novo Detection of Somatic Mutations in High-Throughput Single-Cell Profiling Data Sets.” Nature Biotechnology, July. 10.1038/s41587-023-01863-z.

Pagel, M. 1999. “Inferring the Historical Patterns of Biological Evolution.” Nature 401 (6756): 877–84.

Paterno, Gustavo B., Caterina Penone, and Gijsbert D. A. Werner. 2018. “sensiPhy: An R-package for Sensitivity Analysis in Phylogenetic Comparative Methods.” Methods in Ecology and Evolution / British Ecological Society 9 (6): 1461–67.

Poplin, Ryan, Valentin Ruano-Rubio, Mark A. DePristo, Tim J. Fennell, Mauricio O. Carneiro, Geraldine A. Van der Auwera, David E. Kling, et al. 2017. “Scaling Accurate Genetic Variant Discovery to Tens of Thousands of Samples.” bioRxiv. bioRxiv. 10.1101/201178.

Quinones-Valdez, Giovanni, Ting Fu, Tracey W. Chan, and Xinshu Xiao. 2022. “scAllele: A Versatile Tool for the Detection and Analysis of Variants in scRNA-Seq.” Science Advances 8 (35): eabn6398.

Revell, Liam J., Luke J. Harmon, and David C. Collar. 2008. “Phylogenetic Signal, Evolutionary Process, and Rate.” Systematic Biology 57 (4): 591–601.

Schiffman, Joshua S., Andrew R. D’Avino, Tamara Prieto, Yakun Pang, Yilin Fan, Srinivas Rajagopalan, Catherine Potenski, et al. 2022. “Defining Ancestry, Heritability and Plasticity of Cellular Phenotypes in Somatic Evolution.” bioRxiv. 10.1101/2022.12.28.522128.

Smith, Brian Tilston, William M. Mauck, Brett W. Benz, and Michael J. Andersen. 2020. “Uneven Missing Data Skew Phylogenomic Relationships within the Lories and Lorikeets.” Genome Biology and Evolution 12 (7): 1131–47.

Wang, Kai, Mingyao Li, and Hakon Hakonarson. 2010. “ANNOVAR: Functional Annotation of Genetic Variants from High-Throughput Sequencing Data.” Nucleic Acids Research 38 (16): e164.

Wu, Tianzhi, Erqiang Hu, Shuangbin Xu, Meijun Chen, Pingfan Guo, Zehan Dai, Tingze Feng, et al. 2021. “clusterProfiler 4.0: A Universal Enrichment Tool for Interpreting Omics Data.” Innovation (Cambridge (Mass.)) 2 (3): 100141.

Zafar, Hamim, Yong Wang, Luay Nakhleh, Nicholas Navin, and Ken Chen. 2016. “Monovar: Single-Nucleotide Variant Detection in Single Cells.” Nature Methods 13 (6): 505–7.

Ziegenhain, Christoph, Beate Vieth, Swati Parekh, Björn Reinius, Amy Guillaumet-Adkins, Martha Smets, Heinrich Leonhardt, Holger Heyn, Ines Hellmann, and Wolfgang Enard. 2017. “Comparative Analysis of Single-Cell RNA Sequencing Methods.” Molecular Cell 65 (4): 631–43.e4.

