## Supplementary Information for "Unraveling the phylogenetic signal of gene expression from single-cell RNA-seq data"

a. CINBIO, Universidade de Vigo, 36310 Vigo, Spain
b. Galicia Sur Health Research Institute (IIS Galicia Sur), SERGAS-UVIGO, Spain
C. Department of Biochemistry, Genetics, and Immunology, Universidade de Vigo, 36310 Vigo, Spain

+ Corresponding authors



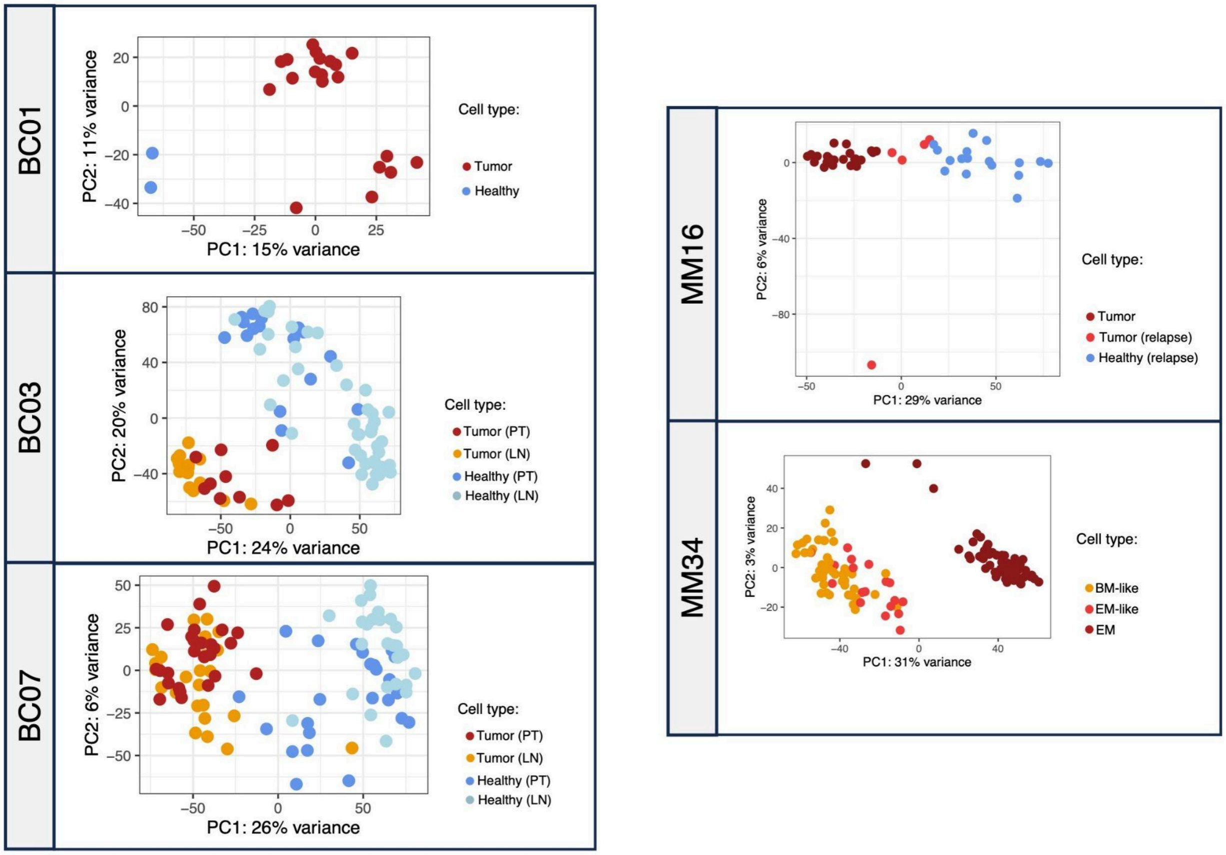


**Figure S1. Gene expression profiles from scRNA-seq data.** Principal component analysis summarizing the gene expression profiles for cells used in this study. Facets represent the different patients. Each cell is depicted by a solid circle with distinct colors highlighting the cell type.

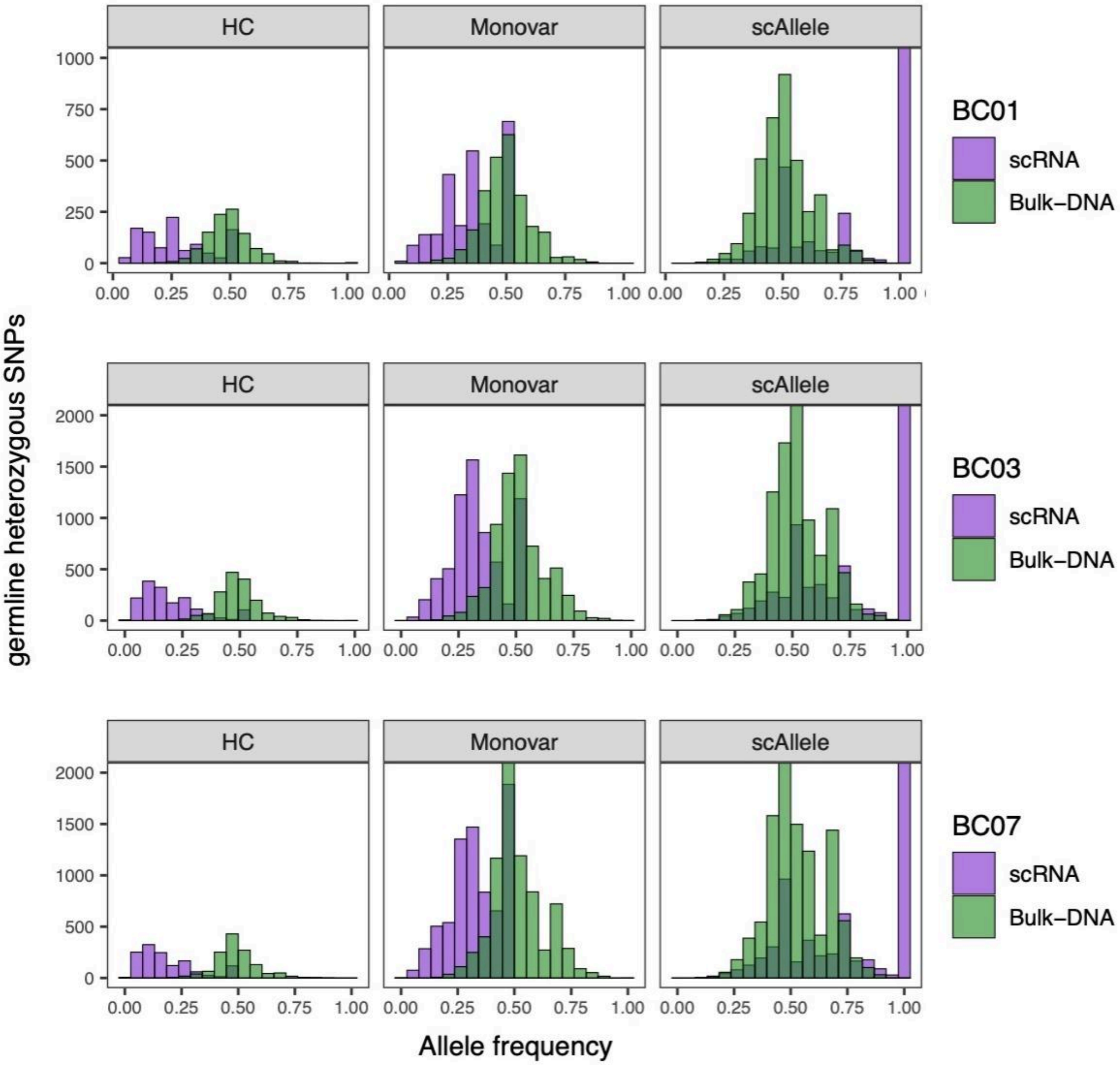


**Figure S2. Variant allele frequency distributions for hetSNPs in bulk DNA-seq and scRNA-seq datasets from BC patients.** Histograms depict the variant allele frequency distribution of hetSNPs in healthy bulk WES (green) and scRNA-seq (purple) data. The allele frequency distribution from WES data was obtained using the allele counts, while for the scRNA-seq data, we used the genotype frequencies in the single-cell population. HC = HaplotypeCaller.

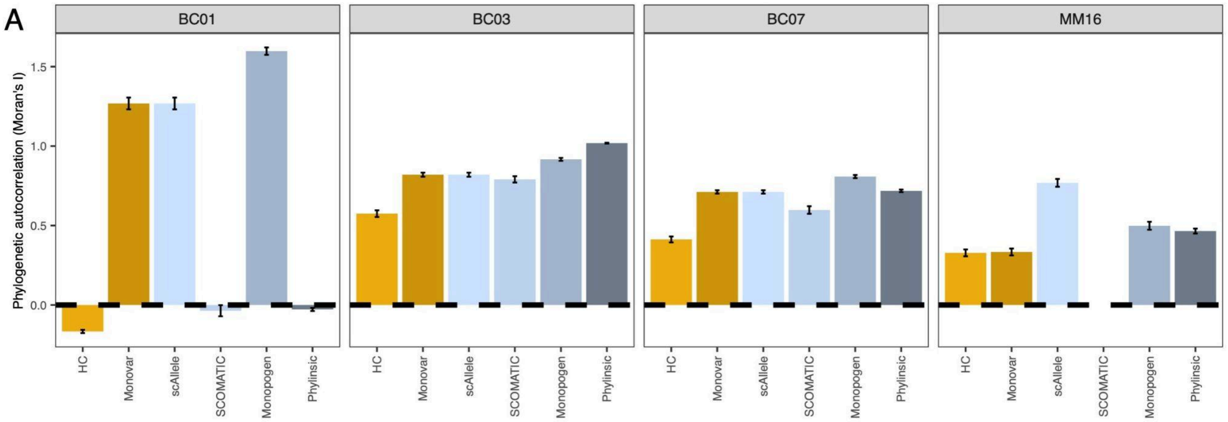


**Figure S3. Phylogenetic autocorrelation.** Bar plots of the tumor vs. healthy Moran's I for the cell phylogenies estimates for the different patients, across bootstrapped trees. Error bars correspond to the standard error of the mean. HC = HaplotypeCaller.

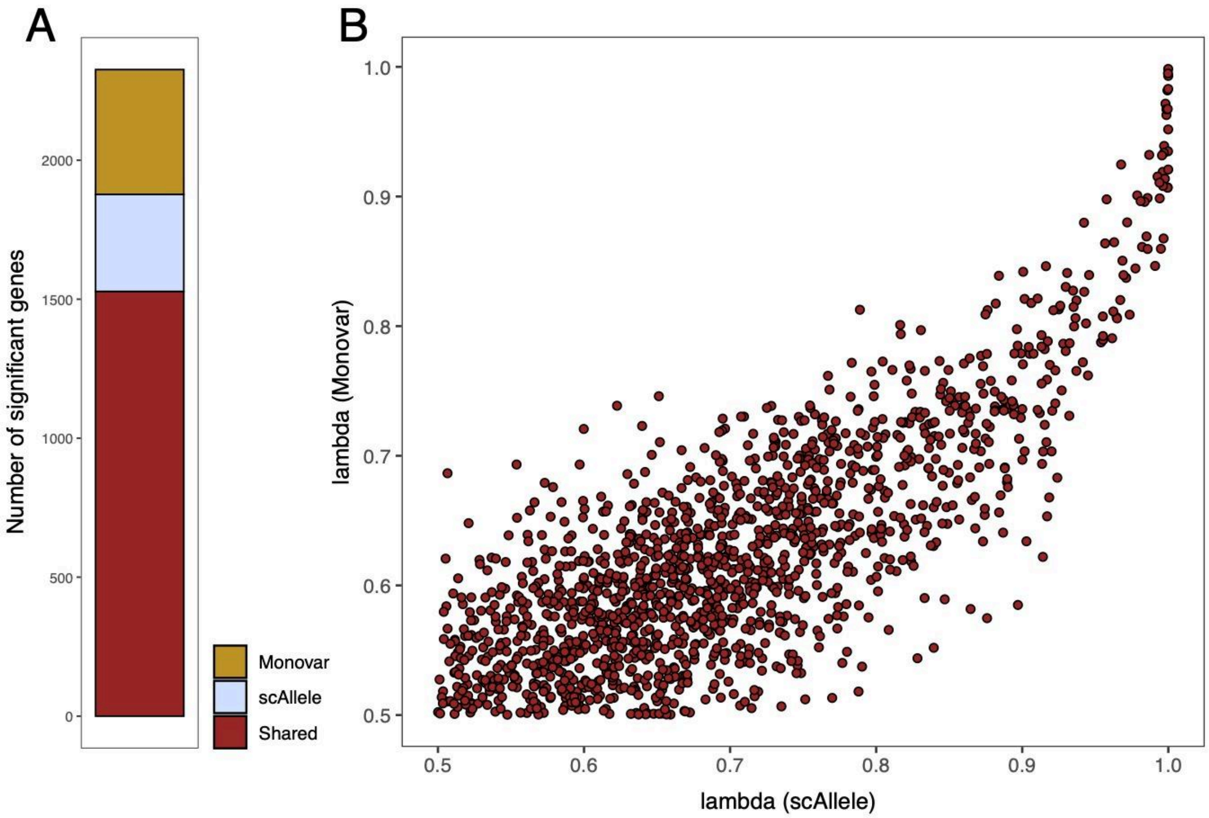


**Figure S4. Phylogenetic signal for scAllele and Monovar in patient MM34. A.** Number of genes showing moderate to high phylogenetic signal (i.e., mean lambda scores ≥ 0.5 and p-values < 0.05) using scAllele and Monovar genotypes. Genes present in both sets are colored in dark red, scAllele-specific genes are colored in light blue, and Monovar-specific genes are colored in gold. **B.** mean lambda scores of shared genes obtained using scAllele and Monovar genotypes.

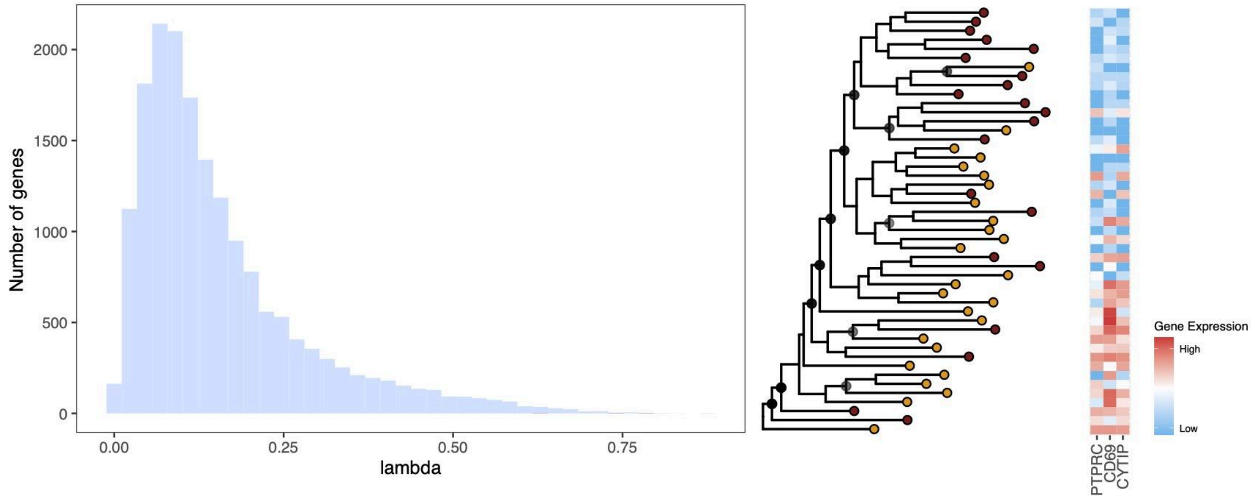


**Figure S5. Phylogenetic signal in BC07. A.** Mean Pagel's lambda score for each gene. Genes showing moderate to high phylogenetic signal (i.e., mean lambda scores ≥ 0.5 and p-values < 0.05) are colored in red. **B.** BC07 phylogenetic tree of tumor cells using the scAllele genotypes. Colors at tips represent sampling locations: primary tumor cells - dark red; lymph-node tumor cells - dark orange. Only bootstrap support values above 50% are shown. Bootstrap scores are depicted using a continuous transparency scale (with solid black circles representing a branch support of 100%). Tile-plot at the right highlights the cell-level gene expression profiles for the three genes with a significant phylogenetic signal. Cells are ordered according to the phylogenetic tree. Gene IDs are shown at the bottom.

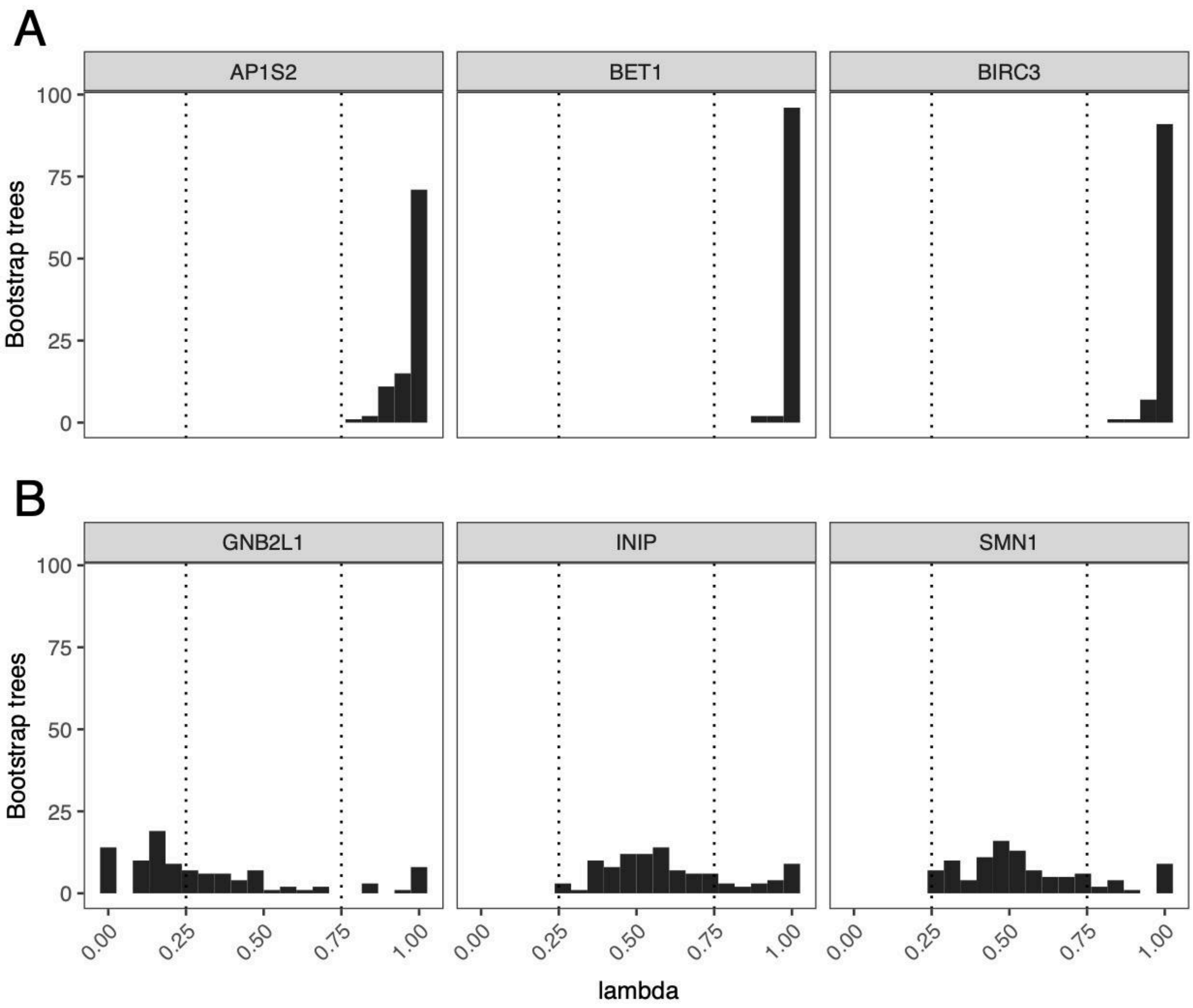


**Figure S6. Phylogenetic signal uncertainty.** Histograms depict lambda scores across 100 bootstrap MM34 trees for six genes, according to the genotypes provided by scAlelle. **A.** Three selected genes with large lambda scores across bootstrap replicates. **B.** Three selected genes with varying lambda scores across bootstrap replicates.
