## Supplementary material for "Unraveling the phylogenetic signal of gene expression from single-cell RNA-seq data": Table S1

| Dataset | SampleID | SRA ID | Cell Type | Sample Type |
|---|---|---|---|---|
| BC01 | BC01_02 | SRR2973279 | tumor | scRNA |
| | BC01_03 | SRR2973280 | tumor | scRNA |
| | BC01_04 | SRR2973281 | tumor | scRNA |
| | BC01_05 | SRR2973282 | tumor | scRNA |
| | BC01_06 | SRR2973283 | tumor | scRNA |
| | BC01_08 | SRR2973284 | tumor | scRNA |
| | BC01_10 | SRR2973285 | tumor | scRNA |
| | BC01_12 | SRR2973286 | tumor | scRNA |
| | BC01_33 | SRR2973287 | tumor | scRNA |
| | BC01_34 | SRR2973288 | tumor | scRNA |
| | BC01_53 | SRR2973289 | tumor | scRNA |
| | BC01_55 | SRR2973290 | tumor | scRNA |
| | BC01_57 | SRR2973291 | tumor | scRNA |
| | BC01_66 | SRR2973292 | tumor | scRNA |
| | BC01_69 | SRR2973293 | tumor | scRNA |
| | BC01_70 | SRR2973294 | tumor | scRNA |
| | BC01_72 | SRR2973295 | tumor | scRNA |
| | BC01_74 | SRR2973296 | healthy | scRNA |
| | BC01_77 | SRR2973297 | tumor | scRNA |
| | BC01_87 | SRR2973298 | tumor | scRNA |
| | BC01_95 | SRR2973299 | tumor | scRNA |
| | BC01_50 | SRR5023384 | healthy | scRNA |
| | BC03LN_01 | SRR2973384 | healthy (LN) | scRNA |
| | BC03LN_04 | SRR2973385 | healthy (LN) | scRNA |
| | BC03LN_05 | SRR2973386 | tumor (LN) | scRNA |
| | BC03LN_06 | SRR2973387 | healthy (LN) | scRNA |
| | BC03LN_10 | SRR2973388 | healthy (LN) | scRNA |
| | BC03LN_11 | SRR2973389 | healthy (LN) | scRNA |
| | BC03LN_13 | SRR2973390 | healthy (LN) | scRNA |
| | BC03LN_14 | SRR2973391 | healthy (LN) | scRNA |
| | BC03LN_17 | SRR2973392 | healthy (LN) | scRNA |
| | BC03LN_19 | SRR2973393 | tumor (LN) | scRNA |
| | BC03LN_20 | SRR2973394 | healthy (LN) | scRNA |

**Acronyms:**
LN: lymphatic node
PT: primary tumor
BM: bone marrow MM
EM: extramedullar MM

| Dataset | NCBI Omnibus ID |
|---|---|
| BC01 | GSE75688 |
| BC03 | GSE75688 |
| BC07 | GSE75688 |
| MM16 | GSE110499 |
| MM34 | GSE110499 |

| | | | | |
|---|---|---|---|---|
| BC03 | BC03LN_21 | SRR2973395 | healthy (LN) | scRNA |
| | BC03LN_23 | SRR2973396 | tumor (LN) | scRNA |
| | BC03LN_25 | SRR2973397 | tumor (LN) | scRNA |
| | BC03LN_26 | SRR2973398 | healthy (LN) | scRNA |
| | BC03LN_27 | SRR2973399 | healthy (LN) | scRNA |
| | BC03LN_28 | SRR2973400 | tumor (LN) | scRNA |
| | BC03LN_31 | SRR2973401 | healthy (LN) | scRNA |
| | BC03LN_32 | SRR2973402 | healthy (LN) | scRNA |
| | BC03LN_34 | SRR2973403 | healthy (LN) | scRNA |
| | BC03LN_35 | SRR2973404 | tumor (LN) | scRNA |
| | BC03LN_37 | SRR2973405 | healthy (LN) | scRNA |
| | BC03LN_38 | SRR2973406 | healthy (LN) | scRNA |
| | BC03LN_42 | SRR2973407 | healthy (LN) | scRNA |
| | BC03LN_44 | SRR2973408 | healthy (LN) | scRNA |
| | BC03LN_47 | SRR2973409 | tumor (LN) | scRNA |
| | BC03LN_48 | SRR2973410 | healthy (LN) | scRNA |
| | BC03LN_49 | SRR2973411 | healthy (LN) | scRNA |
| | BC03LN_50 | SRR2973412 | healthy (LN) | scRNA |
| | BC03LN_51 | SRR2973413 | healthy (LN) | scRNA |
| | BC03LN_53 | SRR2973414 | healthy (LN) | scRNA |
| | BC03LN_56 | SRR2973415 | healthy (LN) | scRNA |
| | BC03LN_57 | SRR2973416 | tumor (LN) | scRNA |
| | BC03LN_58 | SRR5023446 | healthy (LN) | scRNA |
| | BC03LN_61 | SRR2973417 | tumor (LN) | scRNA |
| | BC03LN_62 | SRR2973418 | healthy (LN) | scRNA |
| | BC03LN_63 | SRR2973419 | healthy (LN) | scRNA |
| | BC03LN_64 | SRR2973420 | healthy (LN) | scRNA |
| | BC03LN_65 | SRR2973421 | healthy (LN) | scRNA |
| | BC03LN_68 | SRR2973422 | healthy (LN) | scRNA |
| | BC03LN_69 | SRR2973423 | healthy (LN) | scRNA |
| | BC03LN_70 | SRR2973424 | healthy (LN) | scRNA |
| | BC03LN_71 | SRR2973425 | healthy (LN) | scRNA |
| | BC03LN_72 | SRR2973426 | healthy (LN) | scRNA |
| | BC03LN_73 | SRR2973427 | healthy (LN) | scRNA |
| | BC03LN_78 | SRR2973429 | healthy (LN) | scRNA |

| | | | | |
|---|---|---|---|---|
| | BC03LN_79 | SRR2973430 | healthy (LN) | scRNA |
| | BC03LN_85 | SRR2973431 | healthy (LN) | scRNA |
| | BC03LN_89 | SRR2973433 | tumor (LN) | scRNA |
| | BC03LN_91 | SRR2973434 | healthy (LN) | scRNA |
| | BC03LN_93 | SRR2973435 | healthy (LN) | scRNA |
| | BC03LN_95 | SRR2973436 | healthy (LN) | scRNA |
| | BC03_03 | SRR2973351 | healthy (PT) | scRNA |
| | BC03_06 | SRR2973352 | healthy (PT) | scRNA |
| | BC03_09 | SRR2973353 | tumor (PT) | scRNA |
| | BC03_11 | SRR2973354 | tumor (PT) | scRNA |
| | BC03_13 | SRR2973355 | tumor (PT) | scRNA |
| | BC03_16 | SRR2973356 | tumor (PT) | scRNA |
| | BC03_17 | SRR2973357 | tumor (PT) | scRNA |
| | BC03_19 | SRR2973358 | tumor (PT) | scRNA |
| | BC03_20 | SRR2973359 | tumor (PT) | scRNA |
| | BC03_21 | SRR2973360 | tumor (PT) | scRNA |
| | BC03_23 | SRR2973361 | tumor (PT) | scRNA |
| | BC03_24 | SRR2973362 | healthy (PT) | scRNA |
| | BC03_25 | SRR5023442 | healthy (PT) | scRNA |
| | BC03_26 | SRR2973363 | tumor (PT) | scRNA |
| | BC03_28 | SRR2973364 | healthy (PT) | scRNA |
| | BC03_29 | SRR2973365 | tumor (PT) | scRNA |
| | BC03_35 | SRR2973366 | tumor (PT) | scRNA |
| | BC03_37 | SRR2973367 | tumor (PT) | scRNA |
| | BC03_41 | SRR2973368 | tumor (PT) | scRNA |
| | BC03_43 | SRR2973369 | tumor (PT) | scRNA |
| | BC03_50 | SRR2973371 | healthy (PT) | scRNA |
| | BC03_53 | SRR5023443 | healthy (PT) | scRNA |
| | BC03_57 | SRR2973373 | healthy (PT) | scRNA |
| | BC03_58 | SRR2973374 | healthy (PT) | scRNA |
| | BC03_60 | SRR2973375 | healthy (PT) | scRNA |
| | BC03_66 | SRR5023444 | healthy (PT) | scRNA |
| | BC03_78 | SRR2973377 | healthy (PT) | scRNA |
| | BC03_81 | SRR2973378 | healthy (PT) | scRNA |
| | BC03_85 | SRR5023445 | healthy (PT) | scRNA |

| | | | | |
|---|---|---|---|---|
| | BC03_86 | SRR2973379 | healthy (PT) | scRNA |
| | BC03_92 | SRR2973381 | healthy (PT) | scRNA |
| | BC03_93 | SRR2973382 | healthy (PT) | scRNA |
| | BC03_94 | SRR2973383 | healthy (PT) | scRNA |
| | BC07LN_02 | SRR2973485 | tumor (LN) | scRNA |
| | BC07LN_03 | SRR2973486 | healthy (LN) | scRNA |
| | BC07LN_04 | SRR2973487 | healthy (LN) | scRNA |
| | BC07LN_05 | SRR2973488 | healthy (LN) | scRNA |
| | BC07LN_06 | SRR2973489 | tumor (LN) | scRNA |
| | BC07LN_10 | SRR2973490 | healthy (LN) | scRNA |
| | BC07LN_13 | SRR2973491 | tumor (LN) | scRNA |
| | BC07LN_15 | SRR2973492 | tumor (LN) | scRNA |
| | BC07LN_16 | SRR2973493 | healthy (LN) | scRNA |
| | BC07LN_18 | SRR2973494 | healthy (LN) | scRNA |
| | BC07LN_19 | SRR2973495 | healthy (LN) | scRNA |
| | BC07LN_22 | SRR2973497 | healthy (LN) | scRNA |
| | BC07LN_23 | SRR2973498 | tumor (LN) | scRNA |
| | BC07LN_26 | SRR2973499 | tumor (LN) | scRNA |
| | BC07LN_27 | SRR2973500 | healthy (LN) | scRNA |
| | BC07LN_29 | SRR2973501 | tumor (LN) | scRNA |
| | BC07LN_30 | SRR5023561 | tumor (LN) | scRNA |
| | BC07LN_31 | SRR2973502 | tumor (LN) | scRNA |
| | BC07LN_34 | SRR2973503 | tumor (LN) | scRNA |
| | BC07LN_35 | SRR2973504 | tumor (LN) | scRNA |
| | BC07LN_36 | SRR2973505 | tumor (LN) | scRNA |
| | BC07LN_40 | SRR2973506 | tumor (LN) | scRNA |
| | BC07LN_41 | SRR2973507 | tumor (LN) | scRNA |
| | BC07LN_42 | SRR2973508 | tumor (LN) | scRNA |
| | BC07LN_49 | SRR2973509 | healthy (LN) | scRNA |
| | BC07LN_50 | SRR2973510 | healthy (LN) | scRNA |
| | BC07LN_51 | SRR2973511 | tumor (LN) | scRNA |
| | BC07LN_52 | SRR2973512 | tumor (LN) | scRNA |
| | BC07LN_53 | SRR2973513 | healthy (LN) | scRNA |
| | BC07LN_54 | SRR2973514 | healthy (LN) | scRNA |

| | | | | |
|---|---|---|---|---|
| BC07 | BC07LN_57 | SRR2973515 | tumor (LN) | scRNA |
| | BC07LN_59 | SRR2973516 | healthy (LN) | scRNA |
| | BC07LN_60 | SRR2973517 | tumor (LN) | scRNA |
| | BC07LN_62 | SRR2973518 | healthy (LN) | scRNA |
| | BC07LN_64 | SRR2973519 | healthy (LN) | scRNA |
| | BC07LN_67 | SRR2973520 | healthy (LN) | scRNA |
| | BC07LN_68 | SRR2973521 | healthy (LN) | scRNA |
| | BC07LN_71 | SRR2973522 | healthy (LN) | scRNA |
| | BC07LN_73 | SRR2973523 | tumor (LN) | scRNA |
| | BC07LN_75 | SRR2973524 | tumor (LN) | scRNA |
| | BC07LN_79 | SRR2973525 | tumor (LN) | scRNA |
| | BC07LN_81 | SRR2973526 | healthy (LN) | scRNA |
| | BC07LN_82 | SRR2973527 | tumor (LN) | scRNA |
| | BC07LN_83 | SRR2973528 | healthy (LN) | scRNA |
| | BC07LN_84 | SRR2973529 | healthy (LN) | scRNA |
| | BC07LN_86 | SRR2973530 | tumor (LN) | scRNA |
| | BC07LN_89 | SRR2973531 | healthy (LN) | scRNA |
| | BC07LN_90 | SRR5023562 | healthy (LN) | scRNA |
| | BC07LN_91 | SRR2973532 | tumor (LN) | scRNA |
| | BC07LN_94 | SRR2973533 | healthy (LN) | scRNA |
| | BC07LN_95 | SRR2973534 | tumor (LN) | scRNA |
| | BC07LN_96 | SRR2973535 | healthy (LN) | scRNA |
| | BC07_01 | SRR2973437 | tumor (PT) | scRNA |
| | BC07_03 | SRR2973438 | healthy (PT) | scRNA |
| | BC07_04 | SRR2973439 | healthy (PT) | scRNA |
| | BC07_06 | SRR2973440 | tumor (PT) | scRNA |
| | BC07_07 | SRR5023558 | healthy (PT) | scRNA |
| | BC07_08 | SRR2973441 | tumor (PT) | scRNA |
| | BC07_09 | SRR2973442 | healthy (PT) | scRNA |
| | BC07_12 | SRR2973443 | tumor (PT) | scRNA |
| | BC07_15 | SRR2973444 | tumor (PT) | scRNA |
| | BC07_16 | SRR2973445 | healthy (PT) | scRNA |
| | BC07_18 | SRR2973446 | tumor (PT) | scRNA |
| | BC07_20 | SRR2973447 | tumor (PT) | scRNA |
| | BC07_23 | SRR2973448 | healthy (PT) | scRNA |

| | | | |
|---|---|---|---|
| BC07_24 | SRR2973449 | tumor (PT) | scRNA |
| BC07_27 | SRR2973450 | tumor (PT) | scRNA |
| BC07_30 | SRR2973451 | healthy (PT) | scRNA |
| BC07_31 | SRR2973452 | tumor (PT) | scRNA |
| BC07_35 | SRR2973453 | tumor (PT) | scRNA |
| BC07_36 | SRR2973454 | tumor (PT) | scRNA |
| BC07_38 | SRR2973455 | tumor (PT) | scRNA |
| BC07_39 | SRR2973456 | tumor (PT) | scRNA |
| BC07_42 | SRR2973457 | healthy (PT) | scRNA |
| BC07_43 | SRR2973458 | tumor (PT) | scRNA |
| BC07_44 | SRR2973459 | healthy (PT) | scRNA |
| BC07_50 | SRR2973460 | healthy (PT) | scRNA |
| BC07_52 | SRR2973461 | healthy (PT) | scRNA |
| BC07_55 | SRR5023559 | healthy (PT) | scRNA |
| BC07_56 | SRR2973462 | healthy (PT) | scRNA |
| BC07_57 | SRR2973463 | tumor (PT) | scRNA |
| BC07_58 | SRR5023560 | healthy (PT) | scRNA |
| BC07_59 | SRR2973464 | tumor (PT) | scRNA |
| BC07_60 | SRR2973465 | healthy (PT) | scRNA |
| BC07_61 | SRR2973466 | healthy (PT) | scRNA |
| BC07_63 | SRR2973467 | healthy (PT) | scRNA |
| BC07_65 | SRR2973468 | tumor (PT) | scRNA |
| BC07_66 | SRR2973469 | tumor (PT) | scRNA |
| BC07_67 | SRR2973470 | tumor (PT) | scRNA |
| BC07_68 | SRR2973471 | healthy (PT) | scRNA |
| BC07_69 | SRR2973472 | healthy (PT) | scRNA |
| BC07_72 | SRR2973473 | tumor (PT) | scRNA |
| BC07_73 | SRR2973474 | tumor (PT) | scRNA |
| BC07_75 | SRR2973475 | tumor (PT) | scRNA |
| BC07_78 | SRR2973476 | tumor (PT) | scRNA |
| BC07_79 | SRR2973477 | healthy (PT) | scRNA |
| BC07_80 | SRR2973478 | healthy (PT) | scRNA |
| BC07_83 | SRR2973479 | healthy (PT) | scRNA |
| BC07_87 | SRR2973480 | tumor (PT) | scRNA |
| BC07_89 | SRR2973481 | tumor (PT) | scRNA |

| | | | | |
|---|---|---|---|---|
| | BC07_93 | SRR2973482 | healthy (PT) | scRNA |
| | BC07_95 | SRR2973483 | healthy (PT) | scRNA |
| **BC (Bulk-seq)** | BC01 (Blood) | SRR3022344 | healthy | bulk DNA |
| | BC03 (Blood) | SRR3023076 | healthy | bulk DNA |
| | BC07 (Blood) | SRR3023081 | healthy | bulk DNA |
| | BC01 (tumor) | SRR3022345 | tumor (PT) | bulk DNA |
| | BC03LN (tumor) | SRR3023078 | tumor (LN) | bulk DNA |
| | BC03 (tumor) | SRR3023080 | tumor (PT) | bulk DNA |
| | BC07LN (tumor) | SRR3023082 | tumor (LN) | bulk DNA |
| | BC07 (tumor) | SRR3023083 | tumor (PT) | bulk DNA |
| **MM16** | MM16_1 | SRR6710256 | tumor | scRNA |
| | MM16_10 | SRR6710257 | tumor | scRNA |
| | MM16_12 | SRR6710258 | tumor | scRNA |
| | MM16_15 | SRR6710259 | tumor | scRNA |
| | MM16_16 | SRR6710260 | tumor | scRNA |
| | MM16_17 | SRR6710261 | tumor | scRNA |
| | MM16_21 | SRR6710262 | tumor | scRNA |
| | MM16_24 | SRR6710263 | tumor | scRNA |
| | MM16_3 | SRR6710264 | tumor | scRNA |
| | MM16_31 | SRR6710265 | tumor | scRNA |
| | MM16_35 | SRR6710266 | tumor | scRNA |
| | MM16_5 | SRR6710267 | tumor | scRNA |
| | MM16_52 | SRR6710268 | tumor | scRNA |
| | MM16_54 | SRR6710269 | tumor | scRNA |
| | MM16_57 | SRR6710270 | tumor | scRNA |
| | MM16_6 | SRR6710271 | tumor | scRNA |
| | MM16_61 | SRR6710272 | tumor | scRNA |
| | MM16_63 | SRR6710273 | tumor | scRNA |
| | MM16_69 | SRR6710274 | tumor | scRNA |
| | MM16_74 | SRR6710275 | tumor | scRNA |
| | MM16_8 | SRR6710276 | tumor | scRNA |
| | MM16_83 | SRR6710277 | tumor | scRNA |
| | MM16_9 | SRR6710278 | tumor | scRNA |

| | | | | |
|---|---|---|---|---|
| M | MM16R_01 | SRR6710279 | healthy (relapse) | scRNA |
| | MM16R_02 | SRR6710280 | healthy (relapse) | scRNA |
| | MM16R_07 | SRR6710281 | healthy (relapse) | scRNA |
| | MM16R_16 | SRR6710282 | tumor (relapse) | scRNA |
| | MM16R_21 | SRR6710283 | tumor (relapse) | scRNA |
| | MM16R_23 | SRR6710284 | tumor (relapse) | scRNA |
| | MM16R_30 | SRR6710285 | healthy (relapse) | scRNA |
| | MM16R_32 | SRR6710286 | healthy (relapse) | scRNA |
| | MM16R_34 | SRR6710287 | tumor (relapse) | scRNA |
| | MM16R_38 | SRR6710288 | healthy (relapse) | scRNA |
| | MM16R_39 | SRR6710289 | tumor (relapse) | scRNA |
| | MM16R_41 | SRR6710290 | healthy (relapse) | scRNA |
| | MM16R_44 | SRR6710291 | healthy (relapse) | scRNA |
| | MM16R_51 | SRR6710292 | healthy (relapse) | scRNA |
| | MM16R_56 | SRR6710293 | healthy (relapse) | scRNA |
| | MM16R_59 | SRR6710294 | healthy (relapse) | scRNA |
| | MM16R_60 | SRR6710295 | healthy (relapse) | scRNA |
| | MM16R_65 | SRR6710296 | healthy (relapse) | scRNA |
| | MM16R_70 | SRR6710297 | healthy (relapse) | scRNA |
| | MM16R_73 | SRR6710298 | healthy (relapse) | scRNA |
| | MM16R_87 | SRR6710299 | healthy (relapse) | scRNA |
| | MM16R_93 | SRR6710300 | healthy (relapse) | scRNA |
| | MM34_1 | SRR6710302 | EM-like | scRNA |
| | MM34_10 | SRR6710303 | EM-like | scRNA |
| | MM34_11 | SRR6710304 | EM-like | scRNA |
| | MM34_12 | SRR6710305 | BM-like | scRNA |
| | MM34_13 | SRR6710306 | EM-like | scRNA |
| | MM34_15 | SRR6710307 | BM-like | scRNA |
| | MM34_17 | SRR6710308 | BM-like | scRNA |
| | MM34_18 | SRR6710309 | BM-like | scRNA |
| | MM34_21 | SRR6710310 | EM-like | scRNA |
| | MM34_22 | SRR6710311 | EM-like | scRNA |
| | MM34_23 | SRR6710312 | BM-like | scRNA |
| | MM34_25 | SRR6710313 | BM-like | scRNA |

| | | | | |
|---|---|---|---|---|
| | MM34_27 | SRR6710314 | BM-like | scRNA |
| | MM34_28 | SRR6710315 | BM-like | scRNA |
| | MM34_3 | SRR6710316 | EM-like | scRNA |
| | MM34_32 | SRR6710317 | BM-like | scRNA |
| | MM34_33 | SRR6710318 | BM-like | scRNA |
| | MM34_34 | SRR6710319 | BM-like | scRNA |
| | MM34_35 | SRR6710320 | EM-like | scRNA |
| | MM34_37 | SRR6710321 | BM-like | scRNA |
| | MM34_4 | SRR6710322 | BM-like | scRNA |
| | MM34_42 | SRR6710323 | BM-like | scRNA |
| | MM34_43 | SRR6710324 | EM-like | scRNA |
| | MM34_45 | SRR6710325 | BM-like | scRNA |
| | MM34_46 | SRR6710326 | BM-like | scRNA |
| | MM34_47 | SRR6710327 | EM-like | scRNA |
| | MM34_49 | SRR6710328 | BM-like | scRNA |
| | MM34_50 | SRR6710329 | BM-like | scRNA |
| | MM34_51 | SRR6710330 | BM-like | scRNA |
| | MM34_52 | SRR6710331 | BM-like | scRNA |
| | MM34_53 | SRR6710332 | BM-like | scRNA |
| | MM34_55 | SRR6710333 | EM-like | scRNA |
| | MM34_56 | SRR6710334 | EM-like | scRNA |
| | MM34_57 | SRR6710335 | EM-like | scRNA |
| | MM34_58 | SRR6710336 | BM-like | scRNA |
| | MM34_59 | SRR6710337 | BM-like | scRNA |
| | MM34_6 | SRR6710338 | BM-like | scRNA |
| | MM34_61 | SRR6710339 | BM-like | scRNA |
| | MM34_62 | SRR6710340 | BM-like | scRNA |
| | MM34_63 | SRR6710341 | BM-like | scRNA |
| | MM34_65 | SRR6710342 | BM-like | scRNA |
| | MM34_66 | SRR6710343 | BM-like | scRNA |
| | MM34_69 | SRR6710344 | BM-like | scRNA |
| | MM34_7 | SRR6710345 | EM-like | scRNA |
| | MM34_70 | SRR6710346 | BM-like | scRNA |
| | MM34_71 | SRR6710347 | EM-like | scRNA |
| | MM34_73 | SRR6710348 | EM-like | scRNA |

| | | | | |
|---|---|---|---|---|
| MM34 | MM34_74 | SRR6710349 | EM-like | scRNA |
| | MM34_75 | SRR6710350 | EM-like | scRNA |
| | MM34_78 | SRR6710351 | BM-like | scRNA |
| | MM34_79 | SRR6710352 | BM-like | scRNA |
| | MM34_8 | SRR6710353 | BM-like | scRNA |
| | MM34_80 | SRR6710354 | BM-like | scRNA |
| | MM34_82 | SRR6710355 | BM-like | scRNA |
| | MM34_83 | SRR6710356 | BM-like | scRNA |
| | MM34_85 | SRR6710357 | BM-like | scRNA |
| | MM34_86 | SRR6710358 | BM-like | scRNA |
| | MM34_87 | SRR6710359 | BM-like | scRNA |
| | MM34_88 | SRR6710360 | BM-like | scRNA |
| | MM34_89 | SRR6710361 | BM-like | scRNA |
| | MM34_91 | SRR6710362 | BM-like | scRNA |
| | MM34_92 | SRR6710363 | BM-like | scRNA |
| | MM34_94 | SRR6710364 | BM-like | scRNA |
| | MM34_95 | SRR6710365 | BM-like | scRNA |
| | MM34_96 | SRR6710366 | BM-like | scRNA |
| | MM34EM_1 | SRR6710367 | EM | scRNA |
| | MM34EM_2 | SRR6710368 | EM | scRNA |
| | MM34EM_3 | SRR6710369 | EM | scRNA |
| | MM34EM_5 | SRR6710370 | EM | scRNA |
| | MM34EM_6 | SRR6710371 | EM | scRNA |
| | MM34EM_7 | SRR6710372 | EM | scRNA |
| | MM34EM_11 | SRR6710373 | EM | scRNA |
| | MM34EM_13 | SRR6710374 | EM | scRNA |
| | MM34EM_14 | SRR6710375 | EM | scRNA |
| | MM34EM_15 | SRR6710376 | EM | scRNA |
| | MM34EM_16 | SRR6710377 | EM | scRNA |
| | MM34EM_17 | SRR6710378 | EM | scRNA |
| | MM34EM_18 | SRR6710379 | EM | scRNA |
| | MM34EM_19 | SRR6710380 | EM | scRNA |
| | MM34EM_20 | SRR6710381 | EM | scRNA |
| | MM34EM_21 | SRR6710382 | EM | scRNA |
| | MM34EM_22 | SRR6710383 | EM | scRNA |

| | | | | |
|---|---|---|---|---|
| | MM34EM_23 | SRR6710384 | EM | scRNA |
| | MM34EM_24 | SRR6710385 | EM | scRNA |
| | MM34EM_25 | SRR6710386 | EM | scRNA |
| | MM34EM_30 | SRR6710387 | EM | scRNA |
| | MM34EM_33 | SRR6710388 | EM | scRNA |
| | MM34EM_34 | SRR6710389 | EM | scRNA |
| | MM34EM_35 | SRR6710390 | EM | scRNA |
| | MM34EM_36 | SRR6710391 | EM | scRNA |
| | MM34EM_37 | SRR6710392 | EM | scRNA |
| | MM34EM_38 | SRR6710393 | EM | scRNA |
| | MM34EM_39 | SRR6710394 | EM | scRNA |
| | MM34EM_41 | SRR6710395 | EM | scRNA |
| | MM34EM_43 | SRR6710396 | EM | scRNA |
| | MM34EM_44 | SRR6710397 | EM | scRNA |
| | MM34EM_46 | SRR6710398 | EM | scRNA |
| | MM34EM_48 | SRR6710399 | EM | scRNA |
| | MM34EM_49 | SRR6710400 | EM | scRNA |
| | MM34EM_50 | SRR6710401 | EM | scRNA |
| | MM34EM_51 | SRR6710402 | EM | scRNA |
| | MM34EM_53 | SRR6710403 | EM | scRNA |
| | MM34EM_54 | SRR6710404 | EM | scRNA |
| | MM34EM_56 | SRR6710405 | EM | scRNA |
| | MM34EM_57 | SRR6710406 | EM | scRNA |
| | MM34EM_58 | SRR6710407 | EM | scRNA |
| | MM34EM_59 | SRR6710408 | EM | scRNA |
| | MM34EM_63 | SRR6710409 | EM | scRNA |
| | MM34EM_64 | SRR6710410 | EM | scRNA |
| | MM34EM_66 | SRR6710411 | EM | scRNA |
| | MM34EM_67 | SRR6710412 | EM | scRNA |
| | MM34EM_69 | SRR6710413 | EM | scRNA |
| | MM34EM_72 | SRR6710414 | EM | scRNA |
| | MM34EM_73 | SRR6710415 | EM | scRNA |
| | MM34EM_74 | SRR6710416 | EM | scRNA |
| | MM34EM_75 | SRR6710417 | EM | scRNA |
| | MM34EM_78 | SRR6710418 | EM | scRNA |

| | | | | |
|---|---|---|---|---|
| | MM34EM_80 | SRR6710419 | EM | scRNA |
| | MM34EM_81 | SRR6710420 | EM | scRNA |
| | MM34EM_82 | SRR6710421 | EM | scRNA |
| | MM34EM_85 | SRR6710422 | EM | scRNA |
| | MM34EM_86 | SRR6710423 | EM | scRNA |
| | MM34EM_87 | SRR6710424 | EM | scRNA |
| | MM34EM_88 | SRR6710425 | EM | scRNA |
| | MM34EM_91 | SRR6710426 | EM | scRNA |
| | MM34EM_93 | SRR6710427 | EM | scRNA |
| | MM34EM_94 | SRR6710428 | EM | scRNA |
